# Reconstructing SNP Allele and Genotype Frequencies from GWAS Summary Statistics

**DOI:** 10.1101/2021.04.02.438281

**Authors:** Zhiyu Yang, Peristera Paschou, Petros Drineas

## Abstract

The emergence of genomewide association studies (GWAS) has led to the creation of large repositories of human genetic variation, creating enormous opportunities for genetic research and worldwide collaboration. Methods that are based on GWAS summary statistics seek to leverage such records, overcoming barriers that often exist in individual-level data access while also offering significant computational savings. Here, we propose a novel framework that can reconstruct allelic and genotypic counts/frequencies for each SNP from case-control GWAS summary statistics. Our framework is simple and efficient without the need of any complicated underlying assumptions. Illustrating the great potential of this framework we also propose three summary-statistics-based applications implemented in a new software package (ReACt): GWAS meta-analysis (with and without sample overlap), case-case GWAS, and, for the first time, group polygenic risk score (PRS) estimation. We evaluate our methods against the current state-of-the-art on both synthetic data and real genotype data and show high performance in power and error control. Our novel group PRS method based on summary statistics could not be achieved prior to our proposed framework. We demonstrate here the potential applications and advantages of this approach. Our work further highlights the great potential of summary-statistics-based methodologies towards elucidating the genetic background of complex disease and opens up new avenues for research.

## 1 Introduction

Genomewide association studies (GWAS) have emerged as a powerful tool, leading to the identification of thousands of common genetic variants that underlie human complex disorders and traits. They also led to the creation of large repositories of human genetic variation creating enormous opportunities for further analysis. However, sharing and transferring of individual-level genotype data is often restricted due to privacy concerns as well as logistical issues. On the other hand, GWAS summary statistics, typically including information such as odds ratio (OR)/effect size (beta), standard error (SE), *p*-values, and case/control sample sizes for each SNP being analyzed, are often readily accessible [1]. The availability of such alternative sources of information has spurred intense interest into the development of methodologies seeking to leverage such records effectively in order to retrieve as much information as possible. Besides overcoming barriers in individual-level data access, summary-statistics-based methods also offer advantages in computational costs, which do not scale as a function of the number of individuals in the study [2].

Summary statistics methodologies have been developed to allow a wide array of statistical analyses, including effect size distribution estimation [3, 4]; GWAS meta-analysis and fine mapping [5, 6, 7, 8, 9]; allele frequency and association statistic imputation [10, 11]; heritability and genetic correlation estimation [12, 13, 14, 15]; case-case GWAS [16]; and polygenic prediction [17, 18, 19]. Many of these methods have to incorporate additional information from publicly available sources, such as linkage disequilibrium (LD) statistics from a reference population [12, 10, 20]. Most of the existing methodologies analyzing GWAS summary statistics use the summary statistics (OR, SE, *p*-value) from the input “as is”, without any attempt to recover underlying genotypes, etc. from the summary statistics. Here, we propose a completely novel and simple framework that requires only the assumption of Hardy-Weinberg Equilibrium (HWE) and can convert the summary statistics information into case/control allelic counts for each SNP. Our proposed reconstruction framework provides a completely novel perspective on existing methods and a powerful alternative to summary-statistics-based methods for fixed effect meta-analysis and cc-GWAS. Furthermore, using our framework, we are able to compute group-wise polygenic risk score (PRS) from summary statistics, which, to the best of our knowledge, was completely impossible prior to our work.

We describe the mathematical foundations of our new framework and its application to fixed effect meta-analysis, cc-GWAS, and group-wise PRS estimation. We demonstrate the performance of the proposed methods using simulated and real data and we compare our approach against current state-of-the-art. Our methods are implemented in a new software package: <u>Re</u>constructing <u>A</u>llelic <u>C</u>oun<u>t</u> (ReACt).

## 2 Results

### 2.1 Mathematical foundations

Our framework is motivated by the fact that using summary test statistics from publicly available GWAS allows us to recover allele counts for both the affected and the alternate allele in cases and controls by solving a system of non-linear equations. Interestingly, this rather straight-forward observation has not been documented in prior work. Additionally, assuming that SNPs included in GWAS studies are in Hardy-Weinberg Equilibrium (HWE), we can also reconstruct the structure of the genotype vectors for publicly available GWAS studies from just summary statistics. We can leverage this information in multiple applications, including: *(i)* the computation of the joint effect of a SNP in a meta-analysis involving multiple studies; *(ii)* to obtain the mean polygenic risk score of cases and controls in a population; and *(iii)* to investigate the genetic differences between traits using a case-case GWAS. All of these can be done using only summary statistics, which circumvents the hassle of individual level data sharing and, as an added bonus, considerably reduces the necessary computational time.

We start by introducing some notation that will be useful in this section. Let *a* and *u* represent affected and unaffected allele counts respectively; let superscripts ^cse^ and ^cnt^ represent cases and controls respectively; and let *OR*, *SE*, and *N* be the odds ratio, standard error, and sample sizes obtained from the summary statistics. Thus, for SNP *i*, 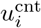 represents the count of the unaffected allele in controls for SNP *i*; similarly, 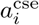 represents the count of the affected allele in cases for SNP *i*; *N* ^cse^ represents the number of cases, etc. We now note that the allelic effect of SNP *i* in case-control GWAS summary statistics can be expressed as follows:

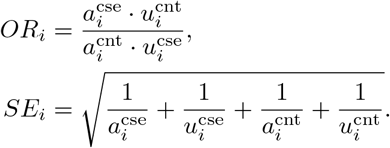

Additionally, sample sizes can be expressed as:

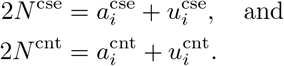

Therefore, solving the system of the above four non-linear equations allows us to recover the allelic counts of SNP *i* for affected and unaffected alleles in cases and controls, by solving for the four unknowns 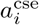, 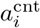, 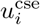, and 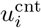. Using these counts, we can trivially obtain allele frequencies in case and control groups and, importantly, by further assuming that the SNPs strictly follow HWE, we can even compute the genotypic counts for each genotype from these frequencies. Note that this reverse engineering scheme applies to GWAS summary statistics generated using a χ^2^ test or logistic regression, but it does not apply to GWAS summary statistics generated by other methodologies. See Section 4.1 and Appendix 6.2 for details.

### 2.2 Fixed effect meta-analysis

#### 2.2.1 Our approach

Armed with allelic and genotypic counts, we can provide a new perspective on fixed-effect GWAS meta-analysis. Instead of the conventional inverse-variance weighted meta-analysis, we can now compute the joint effect of a SNP in a meta-analysis using multiple studies by combining the reconstructed allele and genotype counts from each study and run a *complete* logistic regression on each SNP. Thus, we can essentially proceed with the analysis in exactly the same way as standard GWAS (see Section 4.2 for details). Conceptually, the process is essentially a “mega-analysis” over the combined datasets.

As mentioned in Section 2.1 we can obtain genotypic counts for any SNP over cases and controls from GWAS summary statistics. Then, combining these counts for all available input studies, along with the trait status, we can carry out a logistic regression for this SNP as follows:

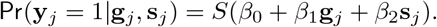

In the above y_*j*_ denotes the binary trait for the *j*th individual, g_*j*_ denotes the respective genotype, and *S*(·) stands for the standard sigmoid function used in logistic regression. Solving for the coefficients *β*_0_, *β*_1_, and *β*_2_ we get the overall SNP effect from the “mega-analysis”. In order to take into account between-study stratification, we introduce an additional variable s_*j*_ as a covariate, using the overall allele frequencies of each study to estimate it. (See Section 4.2 for details.)

#### 2.2.2 Fixed effect meta-analysis: performance evaluation

First, we tested the performance of the proposed fixed-effect meta-analysis approach on synthetic data under various conditions. The simulation was carried out using the Balding-Nichols model, assuming a minor allele frequency of 0.3. For each setting, we predefined the risk for causal SNPs by setting *r* = 1.15/1.2/1.3 as well as the level of population stratification by setting *F*_*st*_ = 0.01/0.05/0.1. Apart from meta-analyzing mutually exclusive datasets, we also tested the performance of our approach under different extents of sample overlap between the input studies: When generating input summary statistics, we evaluated scenarios where the input studies shared *N*_shr_ cases and *N*_shr_ controls, with the value of *N*_shr_ set to zero, 100, and 500 (see Section 4.4.1 for details). We compared power and type I error rates of our approach vs. state-of-the-art tools that are currently widely used for fixed-effect meta-analysis, namely METAL [21] and ASSET [22]. Since the latest stable release of METAL does not include an implementation for sample overlap correction, we used the GitHub version of METAL from [23]. The performance comparison on the meta-analysis of two studies is plotted in Figures 1, 2 and Table S2. Results on synthetic data indicated that our approach has comparable performance with the conventional inverse-variance weighted methods ASSET and METAL, namely

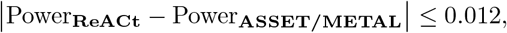

when there is no sample overlap. In scenarios where there were samples shared across input studies, our method (regardless of whether the exact size of the sample overlap is known or is estimated) always showed higher power compared to ASSET, namely

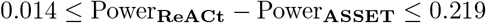

and comparable power to METAL, namely

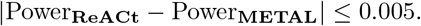

**Figure 1:**
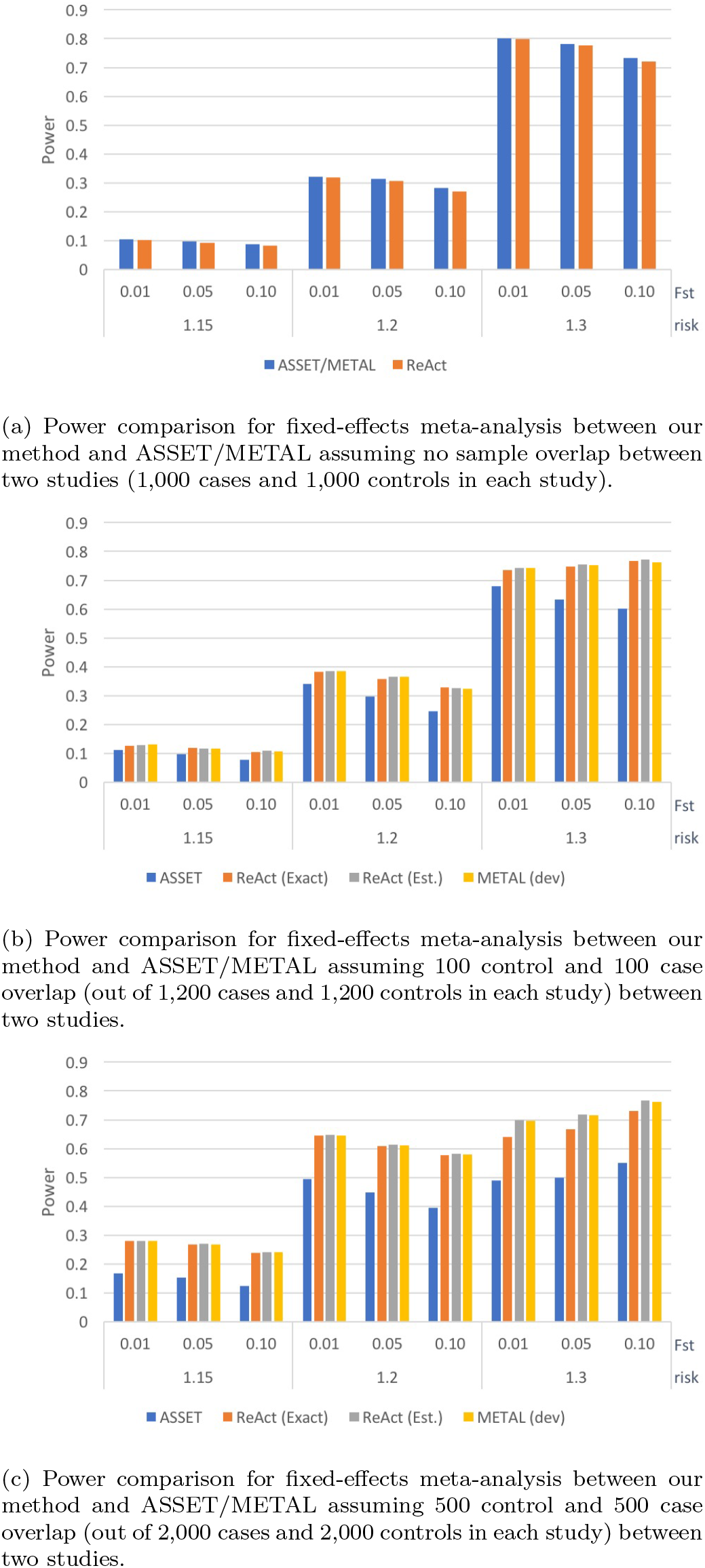
Power of fixed-effect meta-analysis with two input studies under different conditions. We compare the power of our method vs. ASSET/METAL for a significance threshold *p* < 5 · 10^−5^. METAL dev refers to the latest release in GitHub [23]. Two variants of ReACt are tested: Exact and Est, indicating whether the sample overlap was *exactly* known as part of the input or whether it was *estimated*, respectively. Sample overlap indicates the number of cases and controls that were shared between two input studies, ie., a sample overlap equal to 100 means that that there are 100 cases **and** 100 controls shared between two input studies. Total sample sizes for each input study, including the shared samples, are equal to 2,000 when the sample overlap is equal to zero; 2,400 when the sample overlap is equal to 100; and 4,000 when the sample overlap is equal to 500. In each case, the sample is equally split to cases and controls.

**Figure 2:**
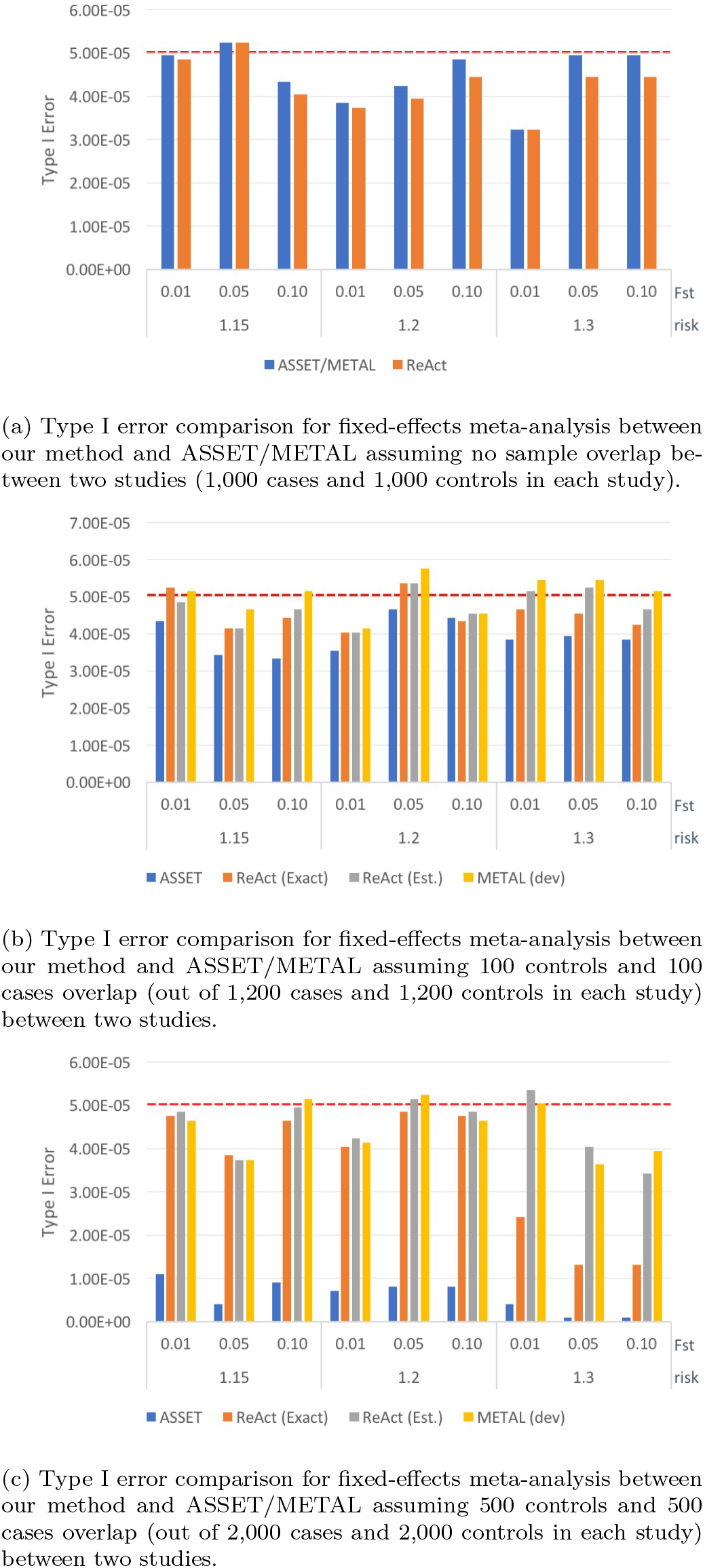
Type I error rate of fixed-effect meta-analysis with two input studies under different conditions. We compared the type I error rate of our method vs. ASSET/METAL for a significance threshold *p* < 5 · 10^−5^. METAL dev refers to the latest release in GitHub [23]. Two variants of ReACt are tested: Exact and Est, indicating whether the sample overlap was *exactly* known as part of the input or whether it was *estimated*, respectively. Sample overlap indicates the number of cases and controls that were shared between two input studies, ie., a sample overlap equal to 100 means that there are 100 cases **and** 100 controls shared between two input studies. Total sample sizes for each input study, including the shared samples, are equal to 2,000 when the sample overlap is equal to zero; 2,400 when the sample overlap is equal to 100; and 4,000 when the sample overlap is equal to 500. In each case, the sample is equally split to cases and controls.

Our advantage in power compared to ASSET was more visible under higher *F_st_* values and larger sample overlaps. In terms of type I error rates, we observed that all methods showed good control on the error rates, while ASSET tended to produce more conservative results. Similar observations can also be made when we meta-analyzed multiple studies; see Table S3 for details.

Beyond power and type I error, we also analyzed the running time of the different methods (see Table S1). Our C implementation of our method in the ReACt software package has not been highly optimized and yet has a running time that is comparable to METAL and is much faster than ASSET. We further tested the performance of our method on real genotype data using a myasthenia gravis dataset from dbGaP (phs000196.v2.p1). The dataset included a total of 964 cases and 1985 controls with 622,328 SNPs after quality control (see Section 4.4.1 for details). In this experiment, we treated the top 13 SNPs with *p*-value stricly less than 10^−5^ from the overall GWAS as “ground truth” and assessed whether various meta-analysis method could pick up these 13 SNPs. Each experiment was carried out over ten iterations: in each iteration, we split the dataset in two equal sized subsets, generated GWAS summary statistics from each of the subsets, and meta-analyzed the resulting summary statistics. We reported average true positive and false positive SNPs counts captured by each method over the ten iterations. Table 1 reports our findings and we note that, perhaps because of the limited power of the dataset or the lack of stratification, the differences in performance were not as visible as what we observed using synthetic data. All methods showed comparable power and type I error. More specifically,

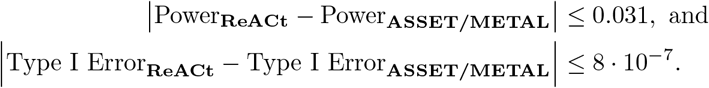

**Table 1:**
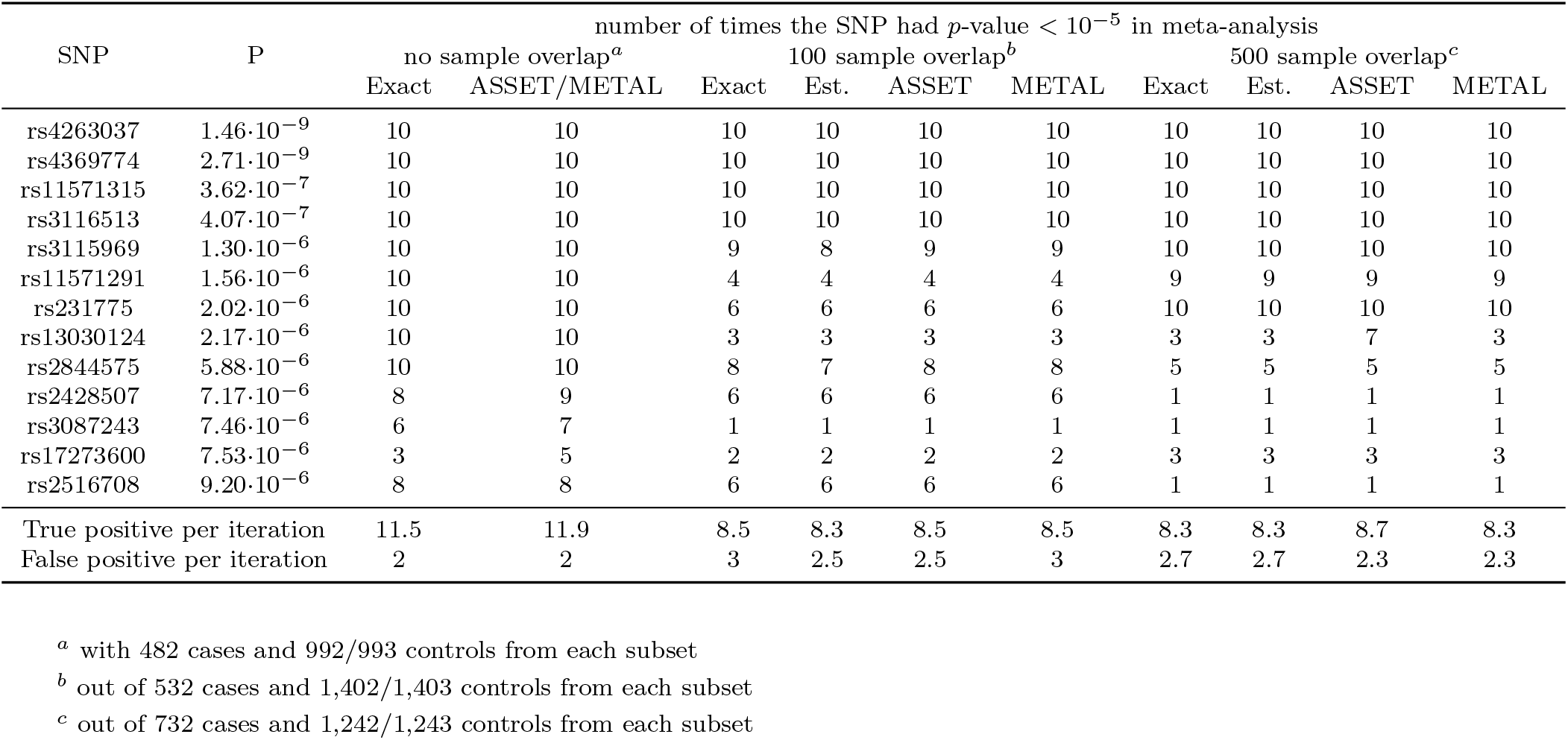
Performance of fixed-effect meta-analysis on real genotype data. We applied our method for fixed-effect meta-analysis to a Myasthenia Gravis GWAS dataset (dbGaP phs000196.v2.p) and compared the performance of our method vs. ASSET/METAL. SNPs with *p*-value strictly less than 10^−5^ in the primary GWAS summary statistics using all samples were treated as “true signals”. In each iteration of an experiment, we split the dataset evenly into two, generated GWAS summary statistics for each subset, and meta-analyzed the summary statistics using our method and ASSET/METAL. We reported the number of times (out of ten iterations) that a “true signal” got captured using the “significance threshold” *p* < 10^−5^ by each method under different sample overlap conditions. METAL dev refers to the latest release in GitHub [23]. Two variants of ReACt are tested: Exact and Est, indicating whether the sample overlap was *exactly* known as part of the input or whether it was *estimated*, respectively. Sample overlap indicates the number of cases and controls that were shared between two input studies, ie., 100 sample overlap means that 100 cases **and** 100 controls were shared between the two studies when the split was carried out. The variable *P* in the table indicates the *p*-value of the target SNP in the primary GWAS using all samples. *True positive per iteration* reports the average number of SNPs with *p*-value strictly less than 10^−5^ in the primary GWAS that were captured in one iteration; and *False positive per iteration* reports the average number of extra SNPs being captured in one iteration.

| SNP | P | number of times the SNP had $p$-value $< 10^{-5}$ in meta-analysis | | | | | | | | | |
| --- | --- | --- | --- | --- | --- | --- | --- | --- | --- | --- | --- |
|  |  | no sample overlap<sup>a</sup> |  | 100 sample overlap<sup>b</sup> |  |  |  | 500 sample overlap<sup>c</sup> |  |  |  |
|  |  | Exact | ASSET/METAL | Exact | Est. | ASSET | METAL | Exact | Est. | ASSET | METAL |
| rs4263037 | $1.46 \cdot 10^{-9}$ | 10 | 10 | 10 | 10 | 10 | 10 | 10 | 10 | 10 | 10 |
| rs4369774 | $2.71 \cdot 10^{-9}$ | 10 | 10 | 10 | 10 | 10 | 10 | 10 | 10 | 10 | 10 |
| rs11571315 | $3.62 \cdot 10^{-7}$ | 10 | 10 | 10 | 10 | 10 | 10 | 10 | 10 | 10 | 10 |
| rs3116513 | $4.07 \cdot 10^{-7}$ | 10 | 10 | 10 | 10 | 10 | 10 | 10 | 10 | 10 | 10 |
| rs3115969 | $1.30 \cdot 10^{-6}$ | 10 | 10 | 9 | 8 | 9 | 9 | 10 | 10 | 10 | 10 |
| rs11571291 | $1.56 \cdot 10^{-6}$ | 10 | 10 | 4 | 4 | 4 | 4 | 9 | 9 | 9 | 9 |
| rs231775 | $2.02 \cdot 10^{-6}$ | 10 | 10 | 6 | 6 | 6 | 6 | 10 | 10 | 10 | 10 |
| rs13030124 | $2.17 \cdot 10^{-6}$ | 10 | 10 | 3 | 3 | 3 | 3 | 3 | 3 | 7 | 3 |
| rs2844575 | $5.88 \cdot 10^{-6}$ | 10 | 10 | 8 | 7 | 8 | 8 | 5 | 5 | 5 | 5 |
| rs2428507 | $7.17 \cdot 10^{-6}$ | 8 | 9 | 6 | 6 | 6 | 6 | 1 | 1 | 1 | 1 |
| rs3087243 | $7.46 \cdot 10^{-6}$ | 6 | 7 | 1 | 1 | 1 | 1 | 1 | 1 | 1 | 1 |
| rs17273600 | $7.53 \cdot 10^{-6}$ | 3 | 5 | 2 | 2 | 2 | 2 | 3 | 3 | 3 | 3 |
| rs2516708 | $9.20 \cdot 10^{-6}$ | 8 | 8 | 6 | 6 | 6 | 6 | 1 | 1 | 1 | 1 |
| True positive per iteration |  | 11.5 | 11.9 | 8.5 | 8.3 | 8.5 | 8.5 | 8.3 | 8.3 | 8.7 | 8.3 |
| False positive per iteration |  | 2 | 2 | 3 | 2.5 | 2.5 | 3 | 2.7 | 2.7 | 2.3 | 2.3 |

<sup>a</sup> with 482 cases and 992/993 controls from each subset
<sup>b</sup> out of 532 cases and 1,402/1,403 controls from each subset
<sup>c</sup> out of 732 cases and 1,242/1,243 controls from each subset

### 2.3 Group PRS

#### 2.3.1 Our approach

Even though we still cannot compute individual level PRS without access to raw genotypes, we observe that, under the additive model, the mean and standard deviation of PRS for a population are just functions of SNP allele frequencies in the target group (see Section 4.3 for details). Therefore, our proposed framework, which returns estimates of allele frequencies for cases and controls using GWAS summary statistics, also allows us to estimate means and standard deviations of PRS for case and control groups using the GWAS summary statistics of the target study. With such information (and a fair assumption of normality in the underlying PRS distribution), we can further run a *t*-test in order to get a *p*-value comparing the difference of PRS between cases and controls.

More specifically, in the additive model, the mean and variance of PRS for a population can be expressed as follows:

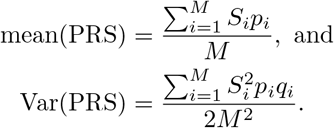

In the above *S*_*i*_ is the weight of SNP *i* inferred from the base summary statistics (typically 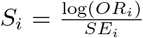), *M* is the total number of SNPs used in the PRS computation, and *p*_*i*_ and *q*_*i*_ = 1 – *p*_*i*_ are allele frequencies of the affected allele and the unaffected allele for SNP *i*. Therefore, we can simply use the allele frequencies of cases and controls that were computed in Section 2.1 in order to get the mean and variance of PRS in cases and controls. See Section 4.3 for details.

#### 2.3.2 Group PRS: Performance evaluation

We first tested our methods on synthetic data without any confounding factors (ie., no stratification). After generating GWAS summary statistics for synthetic base and target datasets, we compared the estimated group means and standard deviations using our method (which operates on summary statistics) with the real group means and standard deviations of PRS computed from the individual level genotypes using PRSice2 [24]. The results successfully proved that in this scenario our method is extremely accurate. See Table 2 which shows typical representative results from our experimental evaluations; essentially identical results were observed in all our experiments on synthetic data.

**Table 2:** Estimated and real group mean and standard deviation of PRS for a synthetic target population. We compared group mean and standard deviation of PRS estimated by REACT from summary statistics of synthetic base and target studies to the real group mean and standard deviation of individual level PRS obtained using summary statistics of the base and individual level genotype of the target computed by PRSice2. Est stands for estimated. Note that the synthetic data is not subject to clumping since the simulation model does not generate LD structure.

| risk | group | Our Method (ReACt) |  | PRSice2 |  |
| --- | --- | --- | --- | --- | --- |
|  |  | est. group mean | est. group sd | real group mean | real group sd |
| 1.15 | cases | 0.0009 | 0.0078 | 0.0009 | 0.0076 |
|  | controls | -0.0037 | 0.0078 | -0.0036 | 0.0081 |
| 1.2 | cases | 0.0016 | 0.0060 | 0.0016 | 0.0059 |
|  | controls | -0.0065 | 0.0060 | -0.0064 | 0.0061 |
| 1.3 | cases | 0.0021 | 0.0041 | 0.0021 | 0.0040 |
|  | controls | -0.0125 | 0.0041 | -0.0125 | 0.0040 |

We further tested our method on real GWAS data, using GWAS summary statistics for myasthenia gravis samples from dbGaP as the base study and assessing its predicting power on 196 *independent* myasthenia gravis cases and 1,057 ancestry-matched controls from [25] for which we had individual level genotypes available. We generated GWAS summary statistics for the base study using standard quality controls and computed GWAS summary statistics for the target dataset as described. We compared the estimated PRS statistics using our methods with the real PRS statistics computed using PRSice2. The results are shown in Table 3; note that since real GWAS datasets are subject to within study population stratification, we did not expect our method to be as accurate as it was on synthetic data without such stratification. There was, however, very high concordance between the results returned by our methods and ground truth. Finally, we applied our methods on summary statistics of eight psychiatric disorders. We evaluated their pairwise PRS predictive power by estimating *t*-test *p*-values. For this experiment, we took into account potential sample overlap between all pairs of base and target studies; see Section 6.3 for details of our sample overlap correction procedure. Results are shown in Table 4 and we observe that, in general, our results coincide with pairwise genetic correlation between disorders as discussed in [7].

**Table 3:**
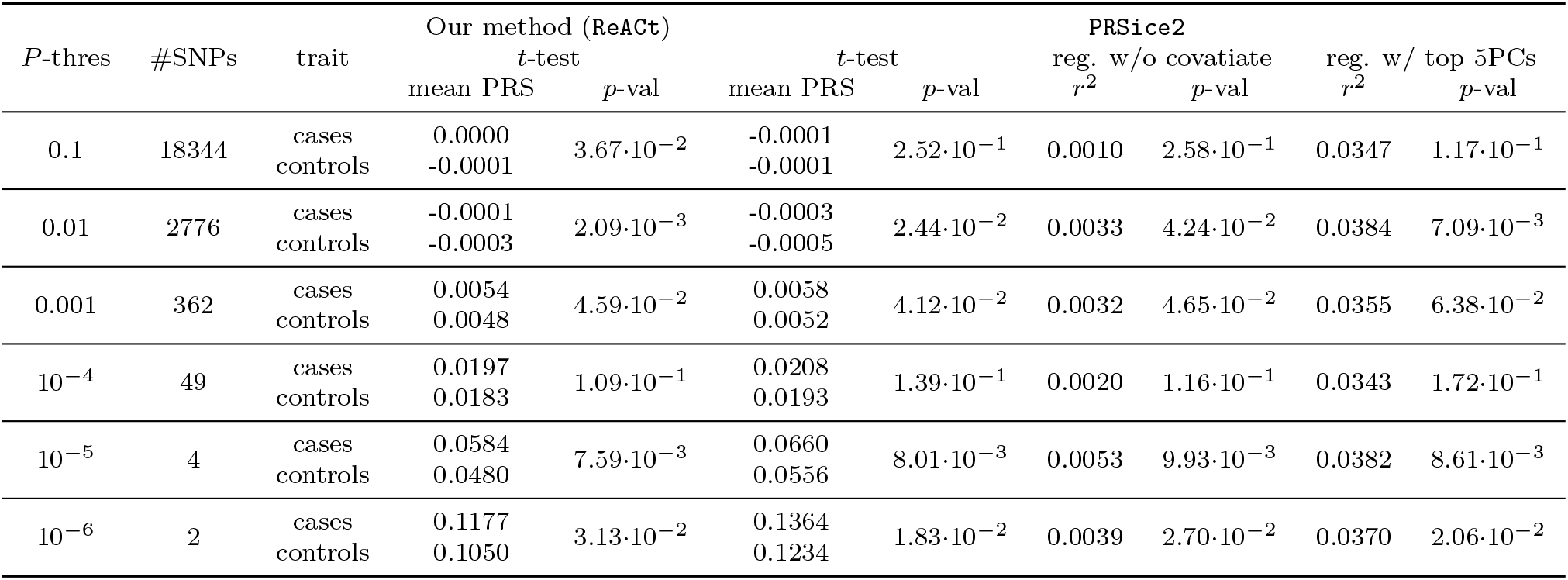
Estimated and real group mean and standard deviation of PRS for a target population of Myasthenia gravis cases and controls. We assessed the performance of our method using a Myasthenia Gravis GWAS dataset (dbGaP phs000196.v2.p) as the base study, and an independent population of 196 Myasthenia Gravis cases and 1,057 ancestry-matched controls as the target population. We generated summary statistics for both base and target populations and estimated group mean PRS and standard deviation of target PRS using ReACt. We computed the individual level PRS for the target study using PRSice2. For both methods, we computed PRS using independent SNPs from the base summary statistics with *p*-values below various thresholds (*P*-thres) and compared the performances under each threshold. For ReACt, mean PRS represents the estimated group mean PRS for cases and controls; *p*-val are the *t*-test *p*-values comparing PRS distribution in cases and in controls. For PRSice2, mean PRS represents real group mean PRS computed from individual level data and *p*-val are the *t*-test *p*-values comparing real PRS distribution in cases and in controls; reg. w/o covariate indicates regression results without covariates, which include the regression *r*^2^ value (reg. *r*^2^) and the *p*-value for the PRS predictor (*p*-val); reg. w/ top 5PCs indicates the regression results including the top five PCs as covariate, , which also included the regression *r*^2^ value (reg. *r*^2^) and the *p*-value for the PRS predictor (*p*-val).

**Table 4:** Using our method to perform PRS comparisons across eight neuropsychiatric disorders. We further applied our method to the summary statistics of eight neuropsychiatric disorders from PGC (see table 12 for details). For each disorder, we used PGC GWAS summary statistics to compute the group mean and standard deviation of PRS for the other seven disorders. All group PRS were estimated using independent SNPs with *p* < 10^−5^in the base summary statistics. We report *p*-values from a *t*-test comparing the group mean PRS of cases against controls in the target study, and cells with deeper blue colors correspond to lower *p*-values. The threshold of significance under multiple testing correction is *p* < 8.93 · 10^−4^.

|  |  | Target |  |  |  |  |  |  |  |
| --- | --- | --- | --- | --- | --- | --- | --- | --- | --- |
|  |  | OCD | TS | ED | ASD | BIP | ADHD | SCZ | MD |
| Base | OCD | - | $5.71 \cdot 10^{-1}$ | $1.26 \cdot 10^{-1}$ | $7.83 \cdot 10^{-2}$ | $9.51 \cdot 10^{-2}$ | $2.64 \cdot 10^{-1}$ | $4.44 \cdot 10^{-1}$ | $6.81 \cdot 10^{-1}$ |
| | TS | $5.17 \cdot 10^{-2}$ | - | $2.31 \cdot 10^{-1}$ | $7.78 \cdot 10^{-1}$ | $3.05 \cdot 10^{-1}$ | $3.57 \cdot 10^{-2}$ | $4.50 \cdot 10^{-1}$ | $5.40 \cdot 10^{-3}$ |
| | ED | $2.95 \cdot 10^{-1}$ | $3.31 \cdot 10^{-1}$ | - | $4.83 \cdot 10^{-1}$ | $4.29 \cdot 10^{-4}$ | $6.28 \cdot 10^{-4}$ | $1.89 \cdot 10^{-2}$ | $3.27 \cdot 10^{-3}$ |
| | ASD | $9.95 \cdot 10^{-1}$ | $7.40 \cdot 10^{-3}$ | $9.00 \cdot 10^{-1}$ | - | $1.77 \cdot 10^{-1}$ | $8.12 \cdot 10^{-4}$ | $1.17 \cdot 10^{-1}$ | $3.98 \cdot 10^{-13}$ |
| | BIP | $3.54 \cdot 10^{-3}$ | $5.82 \cdot 10^{-1}$ | $9.84 \cdot 10^{-13}$ | $4.03 \cdot 10^{-7}$ | - | $1.29 \cdot 10^{-13}$ | $1.08 \cdot 10^{-79}$ | $1.15 \cdot 10^{-19}$ |
| | ADHD | $2.15 \cdot 10^{-1}$ | $1.08 \cdot 10^{-8}$ | $2.32 \cdot 10^{-3}$ | $2.62 \cdot 10^{-45}$ | $9.58 \cdot 10^{-2}$ | - | $1.37 \cdot 10^{-10}$ | $2.88 \cdot 10^{-52}$ |
| | SCZ | $3.23 \cdot 10^{-7}$ | $9.36 \cdot 10^{-1}$ | $4.88 \cdot 10^{-1}$ | $1.28 \cdot 10^{-24}$ | $1.68 \cdot 10^{-133}$ | $2.11 \cdot 10^{-1}$ | - | $7.36 \cdot 10^{-94}$ |
| | MD | $5.09 \cdot 10^{-2}$ | $4.48 \cdot 10^{-1}$ | $3.43 \cdot 10^{-1}$ | $2.08 \cdot 10^{-26}$ | $5.35 \cdot 10^{-9}$ | $6.05 \cdot 10^{-21}$ | $6.10 \cdot 10^{-45}$ | - |

### 2.4 cc-GWAS

#### 2.4.1 Our approach

Similar to our proposed approach for meta-analysis of multiple GWAS datasets using summary statistics, we can also carry out cc-GWAS using regression by simply swapping the labels of the phenotypes. Perhaps the biggest challenge in cc-GWAS is the separation of the differential genetic effects from between-study stratification. To circumvent this issue, we leverage the difference of SNP effects in control groups to estimate the extent of stratification (see Section 4.3.3 for details). Therefore, with a slight modification of the pipeline for meta-analysis of Section 4.2, we introduce an alternate approach for cc-GWAS using our framework.

The underlying theory is quite straightforward and allows us to estimate the genetic differences between two traits of interest using their GWAS summary statistics. Using the genotypic counts we can proceed with logistic regression using only the cases from the two studies:

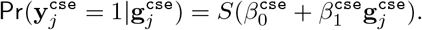

In the above, 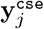 is the binary indicator variable denoting which trait case *j* carries and 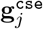 is the genotype of this case. We note that the coefficient 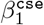 that is part of the output of this regression is a combination of both genetic effects and stratification:

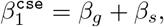

where *β*_*g*_ and *β*_*s*_ are the genetic effect and stratification coefficients. We are only interested in the genetic effect *β*_*g*_ and therefore we need to remove *β*_*s*_. Towards that end, we estimate *β*_*s*_ using the control samples from the input studies; see Section 4.3.3 for details.

#### 2.4.2 CC-GWAS: Performance evaluation

We first tested the performance of our methods on synthetic data. Simulated data were again generated under the Balding-Nichols model, with predefined risks for causal SNPs and the extent of the stratification. Inspired by Peyrot *et al.* [16] we simulated three types of SNPs: *(i)* trait differential SNPs *(ii)* null SNPs; and *(iii)* stress SNPs (see Section 4.4.1 for details). We expect our method to pick up type (i) SNPs and leave the other two. Therefore, in our performance evaluation, we report the power for detecting the type (i) SNPs and type I error rates for picking up type (ii) and (iii) SNPs. Moreover, since we also expect the performance of our method, especially in terms of error control, to vary with sample size, the evaluation was done under different sample sizes in each input study (2,000 cases and 2,000 controls as well as 5,000 cases and 5,000 controls). Power and type I error rates for each type of SNP from the simulation model under each setting are shown in Table 5. The method’s performance was evaluated for *p*-values strictly less than 5 · 10^−5^. For this threshold, our method showed high power and well-controlled type I errors, especially under for lower values of *F*_*st*_. On the other hand, as expected, as stratification increases between two input studies, the power of our method drop and the type I error rates increased for null SNPs. However, as a general trend, we also see a decrease in such error rates when we increase the control sample size. Meanwhile, slightly higher type I error rates for the stress SNPs are observed.

**Table 5:** Performance of cc-GWAS as implemented in ReACt with different sample sizes. Three types of SNPs have been simulated: *(i)* trait differential SNPs; *(ii)* null SNPs; and *(iii)* stress SNPs. . Under each condition, we simulated individual level genotype with these three types of SNPs for *N* cases and *N* controls in each study (*N* = 2, 000 and *N* = 5,000) and generated GWAS summary statistics for each study. and generated GWAS summary statistics for each study respectively. We subsequently used the summary statistics to run cc-GWAS in REACT. We reported the power for detecting type *(i)* SNPs, and false positive rates for picking up type *(ii)* SNPs (Type I err.^(ii)^) and type *(iii)* SNPs (Type I err.^(iii)^) under a significance threshold *p* < 5 · 10^−5^.

| risk | Fst | 2,000 cases, 2,000 controls |  |  | 5,000 cases, 5,000 controls |  |  |
| --- | --- | --- | --- | --- | --- | --- | --- |
|  |  | Power | Type I err.<sup>(ii)</sup> | Type I err.<sup>(iii)</sup> | Power | Type I err.<sup>(ii)</sup> | Type I err.<sup>(iii)</sup> |
| 1.15 | 0.01 | $3.67 \cdot 10^{-2}$ | $2.65 \cdot 10^{-5}$ | $3.16 \cdot 10^{-4}$ | $3.51 \cdot 10^{-1}$ | $1.84 \cdot 10^{-5}$ | $1.87 \cdot 10^{-4}$ |
| | 0.05 | $3.49 \cdot 10^{-2}$ | $9.80 \cdot 10^{-5}$ | $5.26 \cdot 10^{-4}$ | $3.23 \cdot 10^{-1}$ | $6.33 \cdot 10^{-5}$ | $3.58 \cdot 10^{-4}$ |
| | 0.1 | $2.81 \cdot 10^{-2}$ | $2.43 \cdot 10^{-4}$ | $5.02 \cdot 10^{-4}$ | $2.85 \cdot 10^{-1}$ | $1.94 \cdot 10^{-4}$ | $5.21 \cdot 10^{-4}$ |
| 1.2 | 0.01 | $1.54 \cdot 10^{-1}$ | $4.69 \cdot 10^{-5}$ | $2.47 \cdot 10^{-4}$ | $7.16 \cdot 10^{-1}$ | $3.47 \cdot 10^{-5}$ | $2.03 \cdot 10^{-4}$ |
| | 0.05 | $1.34 \cdot 10^{-1}$ | $1.04 \cdot 10^{-4}$ | $5.14 \cdot 10^{-4}$ | $6.62 \cdot 10^{-1}$ | $8.57 \cdot 10^{-5}$ | $3.77 \cdot 10^{-4}$ |
| | 0.1 | $1.23 \cdot 10^{-1}$ | $2.33 \cdot 10^{-4}$ | $5.83 \cdot 10^{-4}$ | $6.03 \cdot 10^{-1}$ | $1.65 \cdot 10^{-4}$ | $5.27 \cdot 10^{-4}$ |
| 1.3 | 0.01 | $5.85 \cdot 10^{-1}$ | $1.63 \cdot 10^{-5}$ | $1.57 \cdot 10^{-4}$ | $9.68 \cdot 10^{-1}$ | $1.43 \cdot 10^{-5}$ | $5.46 \cdot 10^{-4}$ |
| | 0.05 | $5.41 \cdot 10^{-1}$ | $5.31 \cdot 10^{-5}$ | $4.45 \cdot 10^{-4}$ | $9.21 \cdot 10^{-1}$ | $7.35 \cdot 10^{-5}$ | $5.79 \cdot 10^{-4}$ |
| | 0.1 | $4.85 \cdot 10^{-1}$ | $2.63 \cdot 10^{-4}$ | $6.18 \cdot 10^{-4}$ | $8.71 \cdot 10^{-1}$ | $1.67 \cdot 10^{-4}$ | $6.84 \cdot 10^{-4}$ |

Next, we evaluated the performance of our method on real GWAS summary statistics and compared our method with the recently released method of [16]. We analyzed BIP [26] and SCZ [27] datasets, for which case-case GWAS with individual level data was available [28]. We filtered out SNPs that showed untrustworthy estimates of the stratification effect (SE_*s*_ > 0.05, see Section 4.3.3 for details). This reduced our output size from 8,983,436 SNPs being analyzed to 7,110,776 SNPs. Out of those, our analysis revealed a total of 18 genome-wide significant risk loci, including the two regions identified by [28], namely regions 1q25.1 and 20q13.12). We compared our statistics for SNPs that were also analyzed in [16] and results for this comparison are shown in Table 6. The two cc-GWAS methods are mostly comparable. By definition, both we and Peyrot *et al.* [16] only used summary statistics as input, and could not apply the individual level quality control steps of [28]. As a result, both methods identified additional significant loci showing divergent genetic effects between BD and SCZ compared to [28], mainly due to a much larger effective sample size. Results for all genomewide significant risk loci are shown in Table S5.

**Table 6:**
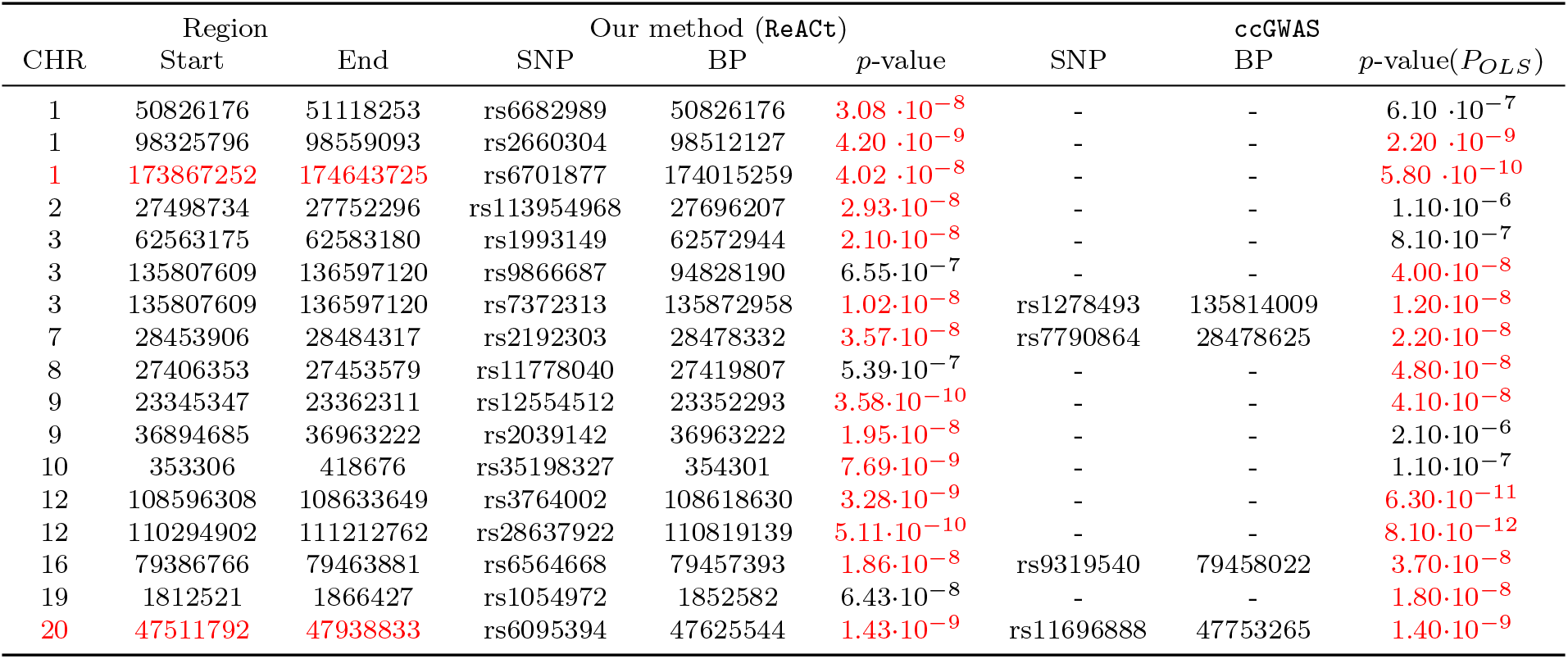
Comparison of genomic regions showing significant divergent genetic effects between BD and SCZ as detected by ReACt and ccGWAS by Peyrot et al [16]. We carried out cc-GWAS with ReACt using summary statistics of BD and SCZ and compared our results with the results from Peyrot et al. Only SNPs that are analyzed in both studies are included for the comparison. Genomic regions that are identified to show significant divergent genetic effects between BD and SCZ in either result are shown. CHR, Start and End are chromosomal and base-pair ranges for the region; SNP, BP and *p*-value (ordinary least squares *p*-values, *P*_*OLS*_, for ccGWAS by Peyrot et al.) are properties of the leading SNP (if the regions is reported genomewide significant) or statistics for the matching SNP (if the region is not reported as genomewide significant, but is detected by the other method); *p*-values in red are leading SNPs that are reported genomewide significant by each method; Regions with CHR, Start and End in red are two loci that were also identified by the case-case GWAS using individual level data [28].

## 3 Discussion

Extracting as much information as possible from easily accessible GWAS summary statistics can help accelerate research that aims to elucidate the genetic background of complex disease, allowing fast sharing of results and datasets while alleviating privacy concerns. Here, we present a simple novel framework to convert SNP statistics from any case-control GWAS back into allelic counts. When summary statistics are generated through simple chi-square tests, the counts will be exact. However, that is not the case for most of the actual GWASs. In practice, this backward reconstruction framework returns “pseudocounts” that correspond to corrected SNP effects after, for example, stratification correction. Therefore, results will not be subject to within-study stratification effects, assuming that the input summary statistics have been generated after stringent quality controls. The framework we propose turns out to be simple, both theoretically and empirically and could broaden the scope of analyses using summary statistics. Not only does it provide new perspectives on some of the existing analytic approaches (meta-analysis and cc-GWAS) but it also expands the potential for novel analyses allowing, for instance, group PRS estimation. We implemented the aforementioned three applications in a readily available software package called ReACt.

As an alternative for fixed-effect meta-analysis, we notice that reconstructing the allelic counts for each SNP allows us to run a full logistic regression model, under the assumption of HWE. The performance of our proposed method turns out to be comparable to conventional approaches while allowing corrections of sample overlaps. Our approach shows increased power in experiments on synthetic data, especially in cases where there is larger *F*_*st*_ difference between the input studies. Our method can therefore be considered as a valid alternative for fixed-effect meta-analysis.

We also propose a novel perspective on case-case association studies (cc-GWAS), allowing an analysis without the need for complicated assumptions or side information apart from sample sizes. To the best of our knowledge, the only publication on summary statistics based case-case GWAS was recently contributed by Peyrot *et al* [16]. Here, we propose a straightforward idea to conduct the case-case GWAS: our approach directly compares the reconstructed allele frequencies of each SNP in two groups of cases, without the requirement to estimate heritabilities or prevalence of disorders as does the method of [16]. Further, we do not need any extra assumptions on the distribution of SNP effects. ReACt analyzes each SNPs independently and, as a result, the analysis is not be subject to any LD structure or number of causal SNPs underlying each disorder. The robustness of our approach is demonstrated by its performance on synthetic data in various scenarios. Similar to the existing cc-GWAS analysis tools [16], `ReACt` showed good control of type I errors in null SNPs (type II SNPs) given sufficiently large control sample sizes for both input studies. It also shows slightly higher, but under-controlled, type I errors in the stress test SNPs (type III SNPs). As pointed out by [16], we do not expect the existence of stress SNPs to be particularly common in practice. We further note that all our experiments on synthetic data were carried out under different levels of population stratification. As expected, our results indicate that the performance of case-case GWAS can be greatly affected by the extent of stratification between the two input studies. We tested the performance of our method for *F*_*st*_ = 0.1, which is a very high end estimate of genetic variation across homo sapiens [29]. Even so, our method still showed good power and type I error rates. For higher confidence in results, we suggest larger sample sizes for both cases and controls, especially when there is higher heterogeneity between the population groups of the two studies. A notable difference between our method and the work of [16] is that we do not filter for SNPs showing association due to differential tagging effects. While analyzing such SNPs, our method behaves more like a direct case-case GWAS using individual level data. Our work is an elegant alternative to [16], offering novel theory and a simple implementation.

Our framework also introduces a novel perspective on case-control PRS. Conventionally, PRS for a target study is only accessible from individual level genotype data. However, even though getting scores for each individual is not feasible, we notice that if we only focus on the differentiation between cases and controls, the group means and standard errors of PRS can in fact be estimated using only summary statistics of both the base and target studies. With such statistics available, a *t*-test can be carried out instead place of logistic regression, which is commonly used for predictability evaluation when the individual level PRS are available. It is worth noting that, for case-control studies, *t*-tests and logistic regression are testing the same hypothesis: whether scores generated from the SNP effect of a base study can differentiate individuals in the target study, or, equivalently, whether the base study can predict the case/control status of samples in the target study. We applied our method to summary statistics of eight psychiatric disorders from PGC for predicting group PRS and found the results in general concordance with the genetic correlation obtained by the work of Lee *et al* [7].

As discussed earlier in our work, our framework is robust against within-study stratification effects, which means that the group means and standard errors returned are corrected for stratification and can be used directly for within-study comparisons. However, we would like to note that the method is still vulnerable to the common weakness of conventional PRS, including differences in population structure between the base and target studies [30]. Users should also keep in mind that general rules of thumb for conventional PRS also apply to our method. For instance, the SNPs used for PRS computations are expected to be independent to a certain extent (clump/prune/LASSO shrink the summary statistics) [19] and as can be observed from the experiments on real data, the predicting power of output PRS will be subject to the power of the base study [31] and the *p*-value threshold chosen by the user. Practices that are not recommended when running conventional PRS (e.g., using results from a GWAS with really small sample size as the base study [31]) are also not recommended in our setting.

We would like to note a couple of potential directions that could further extend our methods. First, the reconstruction scheme that our framework is built upon is based on input summary statistics that are generated using a logistic regression or a *χ*^2^-test. While this is a most common setting, we have not yet explored how to potentially adapt our framework to operate on summary statistics from other models. Also, in this paper, we presented immediate applications of our framework to common tasks in GWAS analyses. An interesting topic for future work would be to incorporate information beyond GWAS summary statistics. For example, one could consider incorporating external information such as LD structure using LD reference maps; such information could for instance be used to attempt to improve the accuracy of sample overlap estimation and extend the group-PRS applications. Furthermore, we could conceivably move towards haplotype reconstruction opening up new possibilities for research.

In conclusion, we introduce a simple and elegant framework that may be used to reconstruct allelic counts and genotypes from GWAS summary statistics. This novel framework highlights the power of summary-statistics-based methodology. We fully expect future extensions will lead to additional applications opening up new possibilities in the quest to identify the genetic background of complex disease.

## 4 Methods

### 4.1 Our framework

#### 4.1.1 Notation

Prior to introducing our methods, we discuss notational conventions. We will reserve the subscript *i* to denote SNP number: given, say, *M* SNPs, *i* will range between one and *M* . Similarly, we will reserve the subscript *ℓ* to denote the study number: given *L* studies from which summary statistics will be meta-analyzed, *ℓ* will range between one and *L*. We assume that all *L* studies released summary statistics on a *common set* of *M* SNPs. For simplicity, we will first describe our methods for the case *L* = 2 (i.e., when exactly two studies are jointly meta-analyzed) and we will generalize our approach in Section 4.2.3 for *L* > 2.

We will use the three-letter shorthand cse for cases and the three-letter shorthand `cnt` for controls. We reserve the variable *a* to represent counts of the affected allele and the variable *u* to represent counts of the unaffected allele. We also reserve the variable *N* to represent counts for the number of cases or controls. Given the above conventions, we now present the following table of allele counts (affected and alternate allele) for SNP *i* (*i* = 1… *M*) in study *ℓ* (*ℓ* = 1… *L*).

Using the above table, we can also compute the frequencies of the affected or alternate allele in cases and controls. Table 8 summarizes frequency notation for SNP *i* (*i* = 1… *M*) in study *ℓ* (*ℓ* = 1… *L*). Obviously,

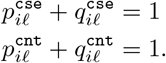

#### 4.1.2 Reconstructing allele counts

Using Table 7, notice that the odds ratio (OR) and its corresponding standard error (SE) for SNP *i* in study *ℓ* are given by the following formulas:

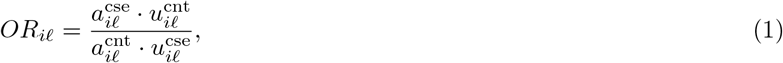

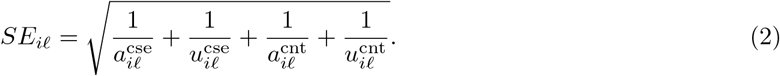

**Table 7:** Table of allele counts for SNP *i* (*i* = 1… *M*) in the *ℓ*-th GWAS (*ℓ* = 1… *L*). The total number of cases for the *ℓ*-th study is 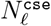 and the total number of controls for the *ℓ*-th study is 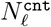. Clearly, the total number of cases and controls in a study is the same for all SNPs, which is why the variable *N* does not depend on *i*. The total number of alleles in cases and controls is equal to twice the number of cases and controls, respectively.

| | $A_1$ (affected allele) | $A_2$ (alternate allele) | Number of alleles |
| --- | --- | --- | --- |
| Cases | $a_{i\ell}^{\text{cse}}$ | $u_{i\ell}^{\text{cse}}$ | $2N_\ell^{\text{cse}}$ |
| Controls | $a_{i\ell}^{\text{cnt}}$ | $u_{i\ell}^{\text{cnt}}$ | $2N_\ell^{\text{cnt}}$ |

**Table 8:** Notations and definitions of (affected or alternate) allele frequencies in cases and controls. The subscripts *i* and *ℓ* indicate SNP number and study number, respectively.

|  |  |
| --- | --- |
| $p_{i\ell}^{\text{cse}} = \frac{a_{i\ell}^{\text{cse}}}{a_{i\ell}^{\text{cse}} + u_{i\ell}^{\text{cse}}}$ | frequency of the *affected allele* $A_1$ in cases |
| $p_{i\ell}^{\text{cnt}} = \frac{a_{i\ell}^{\text{cnt}}}{a_{i\ell}^{\text{cnt}} + u_{i\ell}^{\text{cnt}}}$ | frequency of the *affected allele* $A_1$ in controls |
| $q_{i\ell}^{\text{cse}} = \frac{u_{i\ell}^{\text{cse}}}{a_{i\ell}^{\text{cse}} + u_{i\ell}^{\text{cse}}}$ | frequency of the *alternate allele* $A_2$ in cases |
| $q_{i\ell}^{\text{cnt}} = \frac{u_{i\ell}^{\text{cnt}}}{a_{i\ell}^{\text{cnt}} + u_{i\ell}^{\text{cnt}}}$ | frequency of the *alternate allele* $A_2$ in controls |

Additionally,

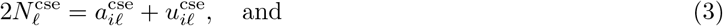

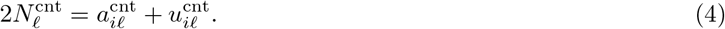

By solving the system of non-linear eqns. (1), (2), (3), and (4), we can recover 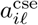, 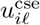, 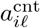, and 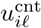 for SNP *i* in study *ℓ*. Notice that *OR*_*iℓ*_, *SE*_*iℓ*_, 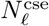, and 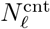 are available from summary statistics. See Appendix 6.2 for details on solving the aforementioned system of non-linear equations.

#### 4.1.3 Reconstructing genotype counts

Given the reconstructed allele counts of Section 4.1.2, we can now reconstruct genotype counts for SNP *i* in the *ℓ*-th study. In order to do this, we need to assume that SNP *i* is in HWE in both case and control groups of study *ℓ*. Note that a well-performed GWAS should have SNPs drastically violating HWE filtered out. More precisely, assume that for SNP *i* in study *ℓ* we have reconstructed its allele table count (Table 7). Then, by assuming that this SNP is in HWE in study *ℓ*, we can compute the number of cases and controls that exhibit a particular genotype. Recall that there are three possible genotypes: *A*_1_*A*_1_, *A*_1_*A*_2_, and *A*_2_*A*_2_. We will represent each genotype by counting the number of copies of the affected allele in each genotype. Thus, *A*_1_*A*_1_ will correspond to two, *A*_1_*A*_2_ will correspond to one, and *A*_2_*A*_2_ will correspond to zero.

Following our notational conventions from Section 4.1.1, we can now compute the entries in Table 9 of genotype counts for SNP *i* in study *ℓ*. It is worth noting that

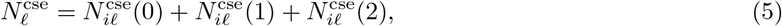

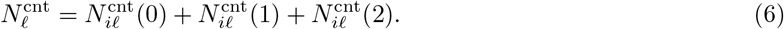

**Table 9:** Genotype counts for cases and controls for SNP *i* in study *ℓ*. Using the above formulas, we can reconstruct the genotype counts for cases and controls for each of the three possible genotypes.

| | $A_1A_1$ (two copies of $A_1$) | $A_1A_2$ (one copy of $A_1$) | $A_2A_2$ (zero copies of $A_1$) |
| --- | --- | --- | --- |
| Cases | $N_{i\ell}^{\text{cse}}(2) = (p_{i\ell}^{\text{cse}})^2 N_{\ell}^{\text{cse}}$ | $N_{i\ell}^{\text{cse}}(1) = 2p_{i\ell}^{\text{cse}} q_{i\ell}^{\text{cse}} N_{\ell}^{\text{cse}}$ | $N_{i\ell}^{\text{cse}}(0) = (q_{i\ell}^{\text{cse}})^2 N_{\ell}^{\text{cse}}$ |
| Controls | $N_{i\ell}^{\text{cnt}}(2) = (p_{i\ell}^{\text{cnt}})^2 N_{\ell}^{\text{cnt}}$ | $N_{i\ell}^{\text{cnt}}(1) = 2p_{i\ell}^{\text{cnt}} q_{i\ell}^{\text{cnt}} N_{\ell}^{\text{cnt}}$ | $N_{i\ell}^{\text{cnt}}(0) = (q_{i\ell}^{\text{cnt}})^2 N_{\ell}^{\text{cnt}}$ |

Next, we reconstruct the genotype vector for SNP *i* in study *ℓ* as follows:

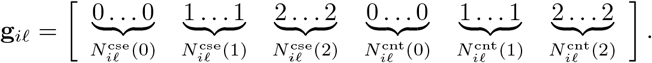

Using eqns. (5) and (6), it is easy to conclude that the vector g_*iℓ*_ has a total of

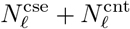

entries, which is equal to the number of samples (cases plus controls) included in the *ℓ*-th study. We can also form the response vector **y**_*ℓ*_ for the *ℓ*-th study, indicating whether a sample is a case (i.e., one) or a control (i.e., zero) as follows:

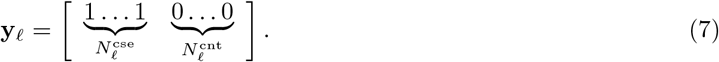

Note that the vectors y_*ℓ*_ and g_*iℓ*_ have the same dimensions (same number of entries). It should be clear that the vector y_*ℓ*_ *is the same for all SNPs* in the *ℓ*-th study and hence does not depend on the SNP number *i*. We conclude the section by discussing the construction of an indicator vector s that will denote the study from which a particular sample in our meta-analysis originated. For the sake of simplicity, assume that we meta-analyze summary statistics from two studies (*L* = 2). Then, following the above discussion, we can construct the genotype vectors g_*i*1_ and g_*i*2_ and concatenate them to construct the overall genotype vector for the *i*-th SNP in both studies:

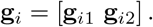

Similarly, we can construct the overall response vector **y** for both studies:

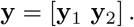

Notice that the vectors **g**_*i*_ and **y** have the same dimensions (number of entries), equal to the number of samples (cases plus controls) in both studies, i.e., equal to

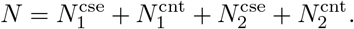

We can now construct the indicator vector **s** as follows:

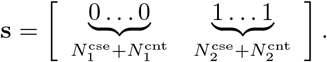

Note that a value of zero in **s** indicates that the corresponding sample belongs to the first study while a value of one in **s** indicates that the corresponding sample belongs to the second study.

### 4.2 Fixed-effect meta-analysis

#### 4.2.1 Logistic regression

We run logistic regression for each SNP separately; recall that we number SNPs in our meta-analysis from one up to *M* . For notational convenience and since we run logistic regression in an identical manner for each SNP, without loss of generality we focus on a single SNP. Let the genotype vector for the selected SNP be denoted by **g**; let s be the study indicator vector; and let **y** be the response vector, as discussed in the previous section. Recall that all three vectors have the same dimensions (same number of entries), equal to *N* , namely the total number of cases and controls in both studies. *Notice that we dropped the subscript i from the vector* **g** *for notational convenience, since our discussion in this section will focus on a fixed SNP i, without loss of generality.*

Using notation from the previous section, while dropping the subscript *i* from the genotype vector **g**, allows us to formulate logistic regression as follows:

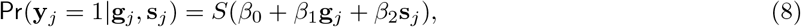

where *S*(*x*) = (1 + *e^−x^*)^−1^ is the sigmoid function; **y**_*j*_ denotes the *j*th entry of the vector **y**; s_*j*_ denotes the *j*th entry of the vector s; and *β*_0_, *β*_1_, and *β*_2_ are the unknown coefficients of the logistic regression formulation. Here *β*_0_ corresponds to the constant offset, *β*_1_ corresponds to the genotype, and *β*_2_ corresponds to the study-of-origin. We also highlight that **g**_*j*_ denotes the *j*th entry of the vector **g**; recall once again that we dropped the subscript *i* from the genotype vector in this section. The range for all subscripts *j* for the above vectors is between one and *N* .

In order to further describe how logistic regression was implemented in our experiments, it will be convenient to introduce additional notation. Let *β* be the vector

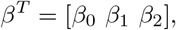

and let **x** be the vector

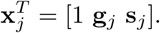

Thus, *β* is the vector of the (unknown) logistic regression coefficients, while 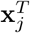 for all *j* = 1… *N* is the vector representing the constant offset, the genotype, and the study origin for the *j*th sample in our meta-analysis. This allows us to rewrite eqn. (8) as follows:

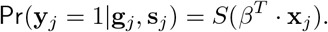

We can now compute the negative log-likelihood (NLL) function for *β* as follows:

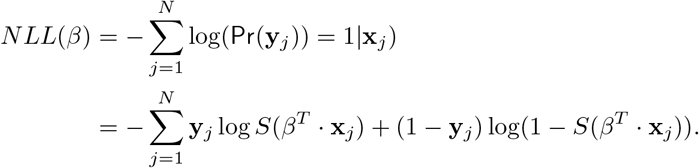

Thus, *β* can be estimated using the Iterative Re-weighted Least Squares (IRLS) algorithm [32] as follows:

**Algorithm 1:**
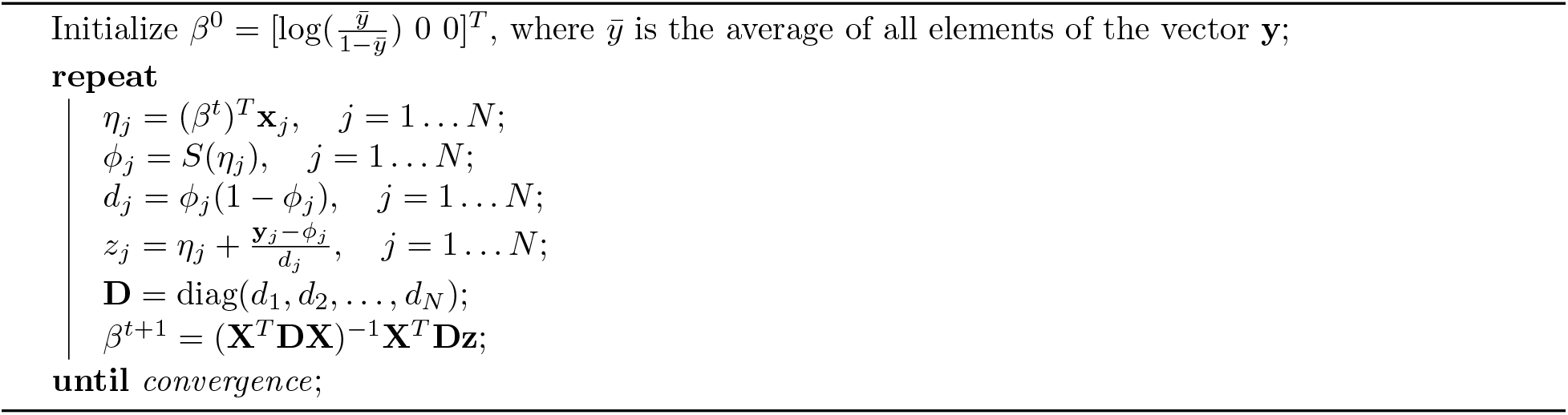
IRLS for maximum likelihood estimate of logistic regression coefficients

In the IRLS algorithm, we let **D** denote the diagonal *N* × *N* matrix whose diagonal entries are *d*_1_, *d*_2_,… . d_*N*_; we let **X** denote the *N* × 3 matrix whose rows are the vectors 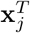 for *j* = 1… *N*; and we let **z** denote the vector whose entries are the *z*_*j*_ for *j* = 1… *N*. Using this notation, the matrix **H** = **X**^*T*^ **DX** is the 3 *×* 3 Hessian matrix of this logistic regression problem. The algorithm iterates over *t* = 0, 1, 2,… and terminates when our convergence criterion, namely the difference ||*β*^t+1^ – *β*^t^ ||_1_ ^1^ drops below the threshold 10^−4^, which is the same threshold as the one used by PLink [33] for logistic regression.

Note that a drawback for logistic regression is that it can produce anti-conservative results under imbalance, which in our case, includes unbalanced sample sizes in cases and controls, as well as unbalanced sample sizes among input studies. We apply Firth bias-corrected logistic regression test [34, 35] to correct for the estimate under input imbalance (triggered when either the total case/control ratio, or maximum/minimum input sample size ratio is greater or equal to 5 by default). This approach has been reported with stable performance in both balanced and unbalanced studies, as well as with rare SNPs [36].

We conclude this section by discussing how to compute a *p*-value for the logistic regression formulation of eqn. (8). First, it is well-known that the standard error for the three coefficients of the logistic regression formulation can be computed by using the inverse of the Hessian matrix **H**. In particular, the standard error for *β*_0_ is equal to 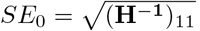; the standard error for *β*_1_ is equal to 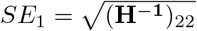; and the standard error for *β*_2_ is equal to 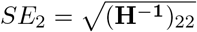. As is typical in association studies, we focus on *SE*_1_, the standard error for the vector of genotypes, and compute the respective *p*-value for the SNP-under-study using the Wald test. More specifically, we find the corresponding *p*-value of a *Z*-distribution for the parameter 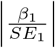.

#### 4.2.2 Correcting for sample overlap (two studies)

Sample overlap between studies can lead to an under-estimation of test statistics variance and results in an inflated test *p*-value. To prevent this from happening, we will use an “effective sample size” correction as follows. Assume that we are given Table 10, which details the number of overlapping samples between the two studies.

**Table 10:** Number of overlapping cases and controls between the two studies. For example, the first cell of the table indicates the number of shared cases between the two studies. In practice, the off-diagonal cells of this table are close to zero, since they indicate cases in one study that became controls in the other study and vice-versa. Large numbers in these off-diagonal cells would indicate high heterogeneity across the two studies, in which case a fixed effect meta-analysis is not recommended.

| Overlapping | Study 2: Case | Study 2: Control |
| --- | --- | --- |
| Study 1: Case | $N_{\text{shr}}^{\text{cse-cse}}$ | $N_{\text{shr}}^{\text{cnt-cse}}$ |
| Study 1: Control | $N_{\text{shr}}^{\text{cse-cnt}}$ | $N_{\text{shr}}^{\text{cnt-cnt}}$ |

Using the counts in Table 10, the number of shared cases between the two studies is equal to:

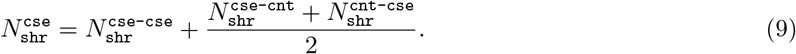

Notice that if the off-diagonal entries in Table 10 are equal to zero then the above number reduces, obviously, to 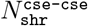. Similarly, we have the number of shared controls equal to:

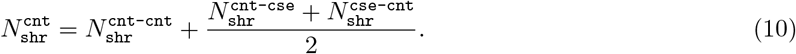

Then, the correction is simply carried out by multiplying the case/control sample size of each input study by a “deflation factor” defined as follows:

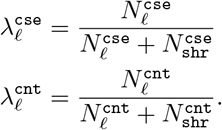

We multiply the sample size for cases (respectively, controls) in each study *ℓ* by 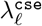 (respectively, 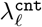) before proceeding with the logistic regression described in Section 4.2.1. See [37] for a similar correction strategy. We finally note that in practice the exact number of overlapping samples between two studies is usually not know. In this case, we followed the approach proposed in [23] to estimate the overlapping sample size.

#### 4.2.3 Meta-analyzing multiple datasets

We now extend our approach to meta-analyze more than two datasets. The main difference with our previously described approach is the handling of the indicator variable for multiple datasets. We can still reconstruct the genotype count for each input study in exactly the same way as in Table 9 as well as the response vector following eqn. (7). Therefore, when multiple studies are meta-analyzed, g_*i*_ and y become

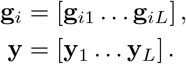

The indicator vector s cannot be binary anymore. Intuitively, one may consider using *L* binary vectors, each to encode samples from each input study. However, this approach would necessitate up to *L*(*L*−1)/2 vectors to encode pairwise sample overlap. This increases the computational complexity by *O*(*L*^2^). A simpler alternative is to use categorical variable as the source study indicator. Note that in this case, different rankings of the studies can lead to completely different results. A straightforward idea is to encode the studies using their population allele frequencies, which can be computed via Table 7 as follows:

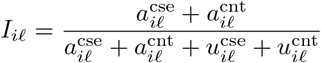

Note this is encoding also controls for population stratification across multiple sample sources. Then, when analyzing *L* studies, the indicator vector **s** becomes:

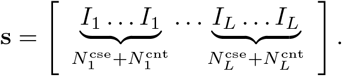

We can now proceed with the logistic regression as in Section 4.2.1. In order to handle sample overlap across multiple studies, we use the subscript 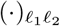 to denote properties of shared samples between two studies *ℓ*_1_ and *ℓ*_2_. Then, generalizing eqns. (9) and (10), we get, for each pair of input studies *ℓ*_1_ and *ℓ*_2_,

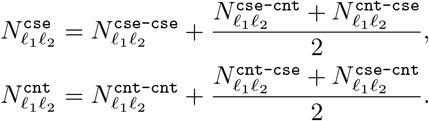

Finally, for any study *ℓ*_1_ = 1… *L*, the sample size correction is

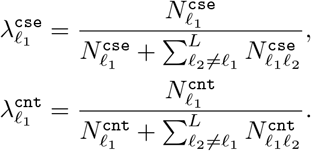

We can now apply 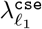 to correct the sample size for cases in study *ℓ*_1_ and we can apply 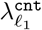 to correct the sample size for controls and proceed with logistic regression.

### 4.3 PRS and cc-GWAS

#### 4.3.1 Mean PRS for cases and controls

Recall that the PRS for the *t*-th individual in the study is computed as:

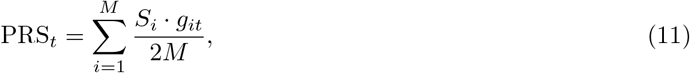

where *g*_*it*_ is the genotype of the *i*-th SNP for the *t*-th individual and *S*_*i*_ is the weight for SNP *i*, which is usually defined as

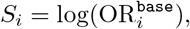

where 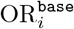 is the odds ratio of SNP *i* in the base summary statistics. Recall from Section 4.1.1 that *M* is the total number of SNPs. Then, in order to compute the average PRS for, say, cases, we simply need to sum up the individual PRS and average over the number of cases. More precisely,

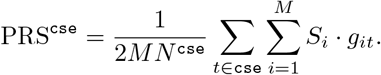

where *N*^cse^ is the number of cases in the target study. The above equation can be rewritten as

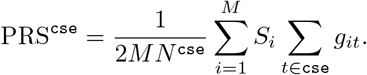

Notice that in an additive model, ∑_*t*∈cse_ *git*/2*N*^cse^ is the allele frequency of SNP *i* over all cases in the target study, which can be computed using only the summary statistics as shown in Section 4.1.3 and Table 8. Thus, the mean PRS under an additive model for cases and controls can be computed as follows:

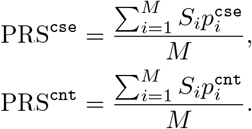

All relevant information for this computation can be easily obtained from the summary statistics of the base and/or target study.

#### 4.3.2 Estimating the standard deviation of the PRS for cases and controls

Interestingly, we can also estimate the standard deviation of the PRS for cases and controls, even Without individual level genotype information, under mild assumptions. First, from eqn. (11), we compute the variance of an individual’s PRS as follows:

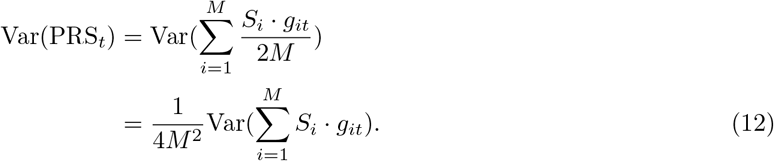

Recall that as a general step prior to the computation of PRS, it is recommended to prune or clump the SNPs used for the PRS computation. Therefore, our first assumption is that the *g*_*it*_’s are pairwise independent. Then, eqn. (12) can be simplified as follows:

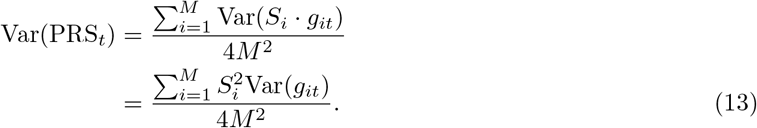

Notice that under an additive model, *g*_*it*_ is a discrete random variable that only takes the value zero, one, and two. Consider all cases and, as in Section 4.1.3 , assume that the SNPs are in HWE. Then, the distribution of *g*_*it*_ in the cases is presented in Table 11. We can now compute the variance of *g*_*it*_ in cases as follows:

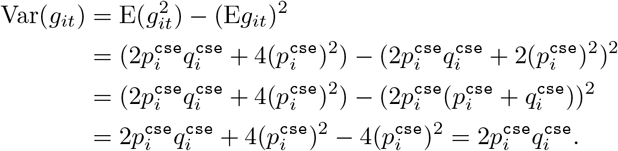

**Table 11:** The probability distribution of *g*_*it*_ for SNP *i*. In this table, 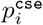 denotes the allele frequency of *A*_1_ in cases and 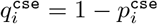.

| $g_{it} = 2$ (two copies of $A_1$) | $g_{it} = 1$ (one copy of $A_1$) | $g_{it} = 0$ (zero copies of $A_1$) |
| --- | --- | --- |
| $(p_i^{\text{cse}})^2$ | $2p_i^{\text{cse}}q_i^{\text{cse}}$ | $(q_i^{\text{cse}})^2$ |

Substituting into eqn. (13), we get

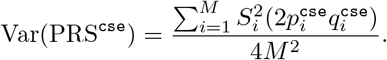

Similarly, we can compute the estimated variance PRS^cnt^ for controls and PRS for the overall population of the target study. To summarize, our estimates are

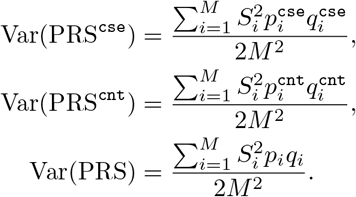

Here *p*_*i*_ is the frequency of allele *A*_1_ for SNP *i* in all samples of the target study, and can be computed as:

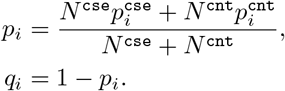

We can now apply a *t*-test in order to obtain a *p*-value for the difference between the PRS distributions in cases and controls. Given the estimated group means and standard deviations for cases and controls, we can further assume that the individual level PRS follow a normal distribution in each group and use the *t*-test statistic as follows:

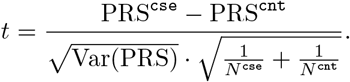

Finally, the degrees of freedom are given by *df* = *N* ^cse^ + *N* ^cnt^ − 2.

#### 4.3.3 cc-GWAS using summary statistics

cc-GWAS is a straight-forward approach to investigate the genetic differences between two traits. However, in practice, it is usually challenging and time consuming, due to restrictions in individual level data sharing. Recently, a method for cc-GWAS that relies only on summary statistics has been proposed in [16]. We propose an alternative perspective on summary-statistics-based cc-GWAS framework, using the foundations of Section 4.1.2.

One of the biggest challenges of cc-GWAS is the differentiation of the genetic effects from trait-trait difference and population stratification. Assume that for a fixed SNP, we run logistic regression focusing only on the cases of the two studies. Let 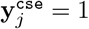 denote that sample *j* is a case from the first study and let 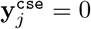 denote that *j* is a case from the second study. Let 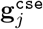 be the genotype of the *j*-th case. Then,

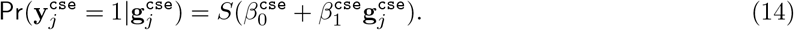

The effect size 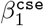 that is the output of logistic regression will include effects from the real genetic differences between trait 1 and trait 2 (*β*_*g*_) as well as from population stratification (*β*_*s*_). We can assume that these two effects are independent of each other:

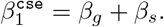

Assume that the control samples from studies one and two *do not carry the traits of interest*. Then, we can estimate the effect of population stratification by running another logistic regression, focusing only on controls from the two studies, as follows:

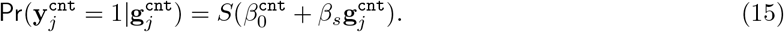

In the above, 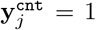 denotes that sample *j* is a control from study one, 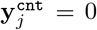 denotes that *j* is a control from study two, and 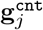 denotes the the genotype for the *j*-th control sample. From this logistic regression, we can get an estimate of the stratification effect *β*_*s*_. Note that along with *β*_*s*_, we will also get a standard error for the estimate of stratification SE_*s*_, which essentially corresponds to the sample size of controls in the two input studies. If we do not have a good amount of controls, SE_*s*_ will turn out to be large, indicating that the estimate for stratification effect is not reliable and the results from the cc-GWAS should be be interpreted carefully.

If SE_*s*_ is small enough, then it is reasonable to assume that the estimate of the stratification effect is credible and we can subsequently treat *β*_*s*_ as a fixed value. Then, the genetic effect from the trait-trait difference that we are interested in is

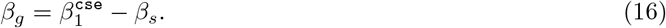

It now follows that the standard error of *β*_*g*_ is

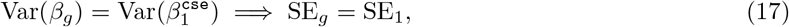

using the derivations of Section 4.1.3. Logistic regressions on cases (eqn. (14)) and controls (eqn. (15)) can be carried out as discussed in Section 4.2.1, with minor changes (include only the designated samples; relabel the dependent variable; and remove the indicator variable). By running these two logistic regressions, we can compute 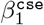, *β*_s_, 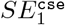, and *SE*_*s*_. Then, using eqns. (16) and (17), we can compute *β*_*g*_ and *SE*_*g*_ for each SNP. Similarly, we can also compute the corresponding *p*-value using a *Z*-distribution for 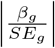.

### 4.4 Experiments

#### 4.4.1 Data

##### Synthetic data

We used the Balding-Nichols model for synthetic genotype generation, assuming a minor allele frequency (MAF) of 0.3 for each SNPs and a relative risk r (*r* = 1.15/1.2/1.3) for the causal SNPs in each population. The simulation was carried out under a range of *F*_*st*_ values (*F*_*st*_ = 0.01/0.05/0.1). For the fixed-effect meta-analysis, we simulated 1,000 cases and 1,000 controls for each input study. A total of 100,000 SNPs were generated, out of which 1,000 are causal SNPs with the predefined risk. Moreover, on top of the independent populations, we also evaluated the performance of ReACt under the presence of sample overlap by introducing a predefined amount of samples shared between each pair of input studies (100 cases, 100 controls overlap; or 500 cases, 500 controls overlap).

For the cc-GWAS, inspired by [16], we used the same simulation model but introduced three types of SNPs for a thorough evaluation of the method’s robustness: *(i)* SNPs with non-zero effect in only one of the studies and zero effect in the other; *(ii)* SNPs with zero effect in both input studies; and *(iii)* SNPs with the same non-zero effect size (predefined *r*) in both input studies. All of the three types of SNPs would suffer from population stratification at a predefined value of *F*_*st*_. In total, 100,000 SNPs were generated, with 1,000 (for each input study) from type (i), 49,000 from type (ii), and 49,000 from type (iii). To investigate the effect of study sizes, we evaluated the method performance on input studies with 2,000 cases and 2,000 controls each, as well as on studies with 5,000 cases and 5,000 controls each.

##### Individual level genotype data

We tested the performance of our fixed-effect meta-analysis method on the myasthenia gravis dataset downloaded from dbGaP (phs000196.v2.p1). This dataset is available as individual level genotypes. We applied basic quality control filters on the dataset, including removing SNPs with a missing rate exceeding 2% or violating the Hardy-Weinberg equilibrium (*p*_*H W E*_ < 0.0001) or having MAF strictly less than 0.05. As a result, 622,328 SNPs and 2,949 samples (964 cases and 1,985 controls) survived and were used for the experiment. For the evaluation of the fixed-effect meta-analysis method, we ran a standard GWAS with all samples and treated SNPs with *p* < 10^−5^ from the results as the “true signals” to be captured. Additionally, to demonstrate the utility of our group PRS method, we used another independent individual level genotype data of cases with myasthenia gravis and matching controls. This dataset has a total sample size of 196 cases and 1,057 controls, with 6,276,739 SNPs included after quality control. This dataset was described in detail in [25].

##### Generating summary statistics

For synthetic data and individual level genotypes, summary statistics were generated using PLink [33], correcting for the top ten principal components (PCs) in the case of admixed datasets. For real individual level genotype data, we divided the samples randomly into two equal sized subsets and ran a GWAS on each subset separately to obtain summary statistics for each subset. We performed ten such random iterations in our experimental evaluations. For the fixed-effect meta-analysis, on top of two independent subsets, we also introduced 100/500 sample overlap to investigate the performance of our methods under more challenging scenarios.

##### Publicly available summary statistics

For group PRS and cc-GWAS, we demonstrated the applicability of our methods using publicly available summary statistics. We chose the summary statistics of eight neuropsychiatric disorders made available by the Psychiatric Genomics Consortium (PGC), since the underlying relationships between this set of disorders has been relatively well-studied. Information on the eight summary statistics can be found in Table 12.

**Table 12:** Information on summary statistics for the eight psychiatric disorders used in the experiments. Note that we used summary statistics only for samples of European ancestry. For MD, we used the summary statistics generated by UK biobank, excluding the 23andMe samples; for BIP, we used the summary statistics including all three patient sub-types.

| Disorder | #Cases | #Controls | Total | #SNPs | Reference |
| --- | --- | --- | --- | --- | --- |
| obsessive-compulsive disorder (OCD) | 2,688 | 7,037 | 9,725 | 8,409,516 | [38] |
| Tourette syndrome (TS) | 4,819 | 9,488 | 14,307 | 8,947,432 | [39] |
| eating disorder (ED) | 3,495 | 10,982 | 14,477 | 10,641,224 | [40] |
| autism spectrum disorder (ASD) | 18,382 | 27,969 | 46,351 | 9,112,386 | [41] |
| bipolar disorder (BIP) | 20,352 | 31,358 | 51,710 | 13,413,244 | [26] |
| schizophrenia (SCZ) | 36,989 | 113,075 | 150,064 | 9,075,843 | [27] |
| attention-deficit/hyperactivity disorder (ADHD) | 19,099 | 34,194 | 53,293 | 8,094,094 | [42] |
| major depression (MD) | 69,232 | 161,009 | 230,241 | 9,874,289 | [43] |

#### 4.4.2 Evaluation metrics

##### Fixed-effect meta-analysis

For synthetic experiments, results after performing the meta-analysis were compared with the predefined causal variants. Power and type I error rate under each experimental condition were reported as an average of ten independent repetitions. For real genotype data, in each iteration, we meta-analyzed summary statistics of two subsets using the proposed methods and standard approaches and compared results with the GWAS results on the complete dataset. We again reported results averaged over ten iterations (random splits) showing, on average, how many times a SNP reported as a “true signal” in the overall GWAS got picked up by each meta-analysis method (true positive) as well as how many extra SNPs each method identified (false positive). The performance on real genotype data was also evaluated under 0/100/500 sample overlap. Sample size for each subset under different conditions was 482 cases, 993 controls with no sample overlap; 532 cases, 1043 controls with 100 cases and 100 controls overlap; and 732 cases, 1243 controls with 500 cases and 500 controls overlap.

We compared the performance of ReACt in terms of accuracy as well as running time with METAL [21] and ASSET [22], which are both widely used tools for fixed-effect meta-analysis. Note that the latest stable release of `METAL` does not have the sample overlap correction functionality implemented. Therefore, for performance comparison, we used the *development version* available on GitHub [23].

##### Group PRS

In order to show that our method outputs reliable estimates of the group-wise statistics for PRS without accessing individual level genotypes, we compared the output of our method to the true group mean and standard deviation computed from the individual level PRS on synthetic data, as described in Section 4.4.1. Performance was evaluated under with a fixed 0.05 *F*_*st*_ between the base and target studies. For a pair of base and target studies, we estimated the mean PRS for case/control groups as well as their standard deviation using SNPs with *p*-values strictly less than 5 · 10^−5^ in the summary statistics. We also computed the individual level PRS using PRSise to obtain the true group mean and standard deviation. Our experiments show that our estimates are numerically close to the real values. Next, we evaluated the performance of ReACt on real GWAS datasets, where the individual level genotype of the target study was available. For this experiment, we used GWAS summary statistics of myasthenia gravis samples from dbgap as the base study (see Section 4.4.1 for details) and an independent group of myasthenia gravis cases and matching controls as the target population [25]. We clumped the base summary statistics using the European samples from 1000 Genome Project as reference, under parameters `--clump-p1 1 --clump-kb 250 --clump-r2 0.1`. We tested the method and reported results under a range of *p*-value thresholds (0.1, 0.01, 0.001, 10^−4^, 10^−5^, and 10^−6^). For each threshold, we used only independent SNPs with a *p*-value smaller than the respective threshold from the base summary statistics for PRS calculation, using both ReACt and PRSice2 [24]. We reported the mean PRS of cases and controls, as well as the resulting *p*-value from *t*-test. In the case of PRSice2, we also reported the regression *r*^2^ value and *p*-value for the PRS predictor with and without correcting for covariates (ie., the top five principal components).

Finally we applied ReACt to summary statistics of eight neuropsychiatric disorders (OCD, TS, ED, ADHD, ASD, BIP, SCZ and MDD, see Section 4.4.1 for details) and reported the pairwise PRS prediction power in terms of *t*-test *p*-values for the difference between case/control group PRS means. Prior to the group PRS computation, each base summary statistics was clumped using PLink [33] using parameters `--clump-p1 1 --clump-kb 250 --clump-r2 0.1`, with the European samples from 1000 Genome Project as a reference. All PRS values were estimated using independent SNPs with *p*-values strictly less than 10^−5^ from the base summary statistics.

##### cc-GWAS

Out of the three types of SNPs generated for the cc-GWAS evaluation (see Section 4.4.1), we expect ReACt to pick up only type (i) SNPs as they have been designed to be the trait differential SNPs. Therefore, we reported the power of ReACt based on the number of type (i) SNPs that were identified as well as type I error rates for type (ii) SNPs and type (iii) SNPs. Since the randomness introduced by the simulation could lead to false positives that were not due to the method itself, we filtered out type (iii) SNPs showing extreme differences in effect size between studies, by removing type (iii) SNPs with |*OR*_*i*1_ − *OR*_*i*2_| ≥ 0.1 from performance evaluation. Here *OR*_*i*1_ corresponds to the odd ratio for the *i*th SNP in the first study and *OR*_*i*2_ corresponds to the odd ratio for the *i*th SNP in the other study. Since all three types of SNPs suffered from population stratification, we evaluated the performance of ReACt under a challenging scenario. Besides simulation, experiments using summary statistics for schizophrenia (SCZ) [44] and bipolar disorder (BIP) [45] were also carried out. These two disorders were chosen due to the existence of case-case association study using the individual level genotypes [28]. We tested ReACt using the summary statistics and compared the results with the existing case-case association study between SCZ and BIP to see whether it could detect possible genetic differences between the two disorders. Since no individual level quality control could be carried out, we expected our results to correspond to a case-case GWAS including 36,989 cases from SCZ and 20,352 cases from all three sub-types of BIP (type 1, type 2, and schizoaffective bipolar disorder). For the analysis, we excluded SNPs on the X-chromosome, MHC region (chr6: 25,000,000 - 35,000,000BP), and the inversion on chromosome 8 (chr8: 7,000,000 - 15,000,000BP). As a result, a total of 8,983,436 SNPs shared between both summary statistics were used for the analysis. The results were compared in detail with the results reported by the cc-GWAS in [16].

## 5 Conclusion

In summary, we propose a simple, novel framework that reconstructs allelic counts of each SNP from the summary statistics of case-control GWAS. Additionally, we evaluate our framework on three applications and provide light and easily modifiable implementations of our methods in the software package ReACt. Given the simplicity of the proposed approach, both theoretically and empirically, we believe that this framework has significant potential for further developments.

## URL

This is a preliminary implementation for ReACt: https://github.com/Paschou-Lab/ReAct Please contact us if you identify any bug when using this version of ReACt and we will keep improving.

## Acknowledgements

The Genome-wide Association Study of Myasthenia Gravis from dbGaP (phs000196.v2.p1) was funded by the Myasthenia Gravis Foundation, a Bequest from Geraldine Weinrib, and the Intramural Research Program of the National Institute on Aging, National Institutes of Health.

This research was funded by NSF grants 1715202 and 2006929.

We would also like to thank our colleague Pritesh Jain for contributing the name ReACt for the tool being developed.

## 6 Supplementary Material

### 6.1 Supplementary tables

**Table S1:**
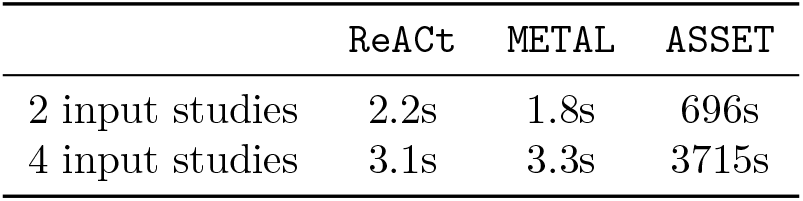
Average running time in seconds for fixed effect meta-analysis for ReACt, METAL, and ASSET. All experiments were performed at Purdue’s Snyder cluster on a dedicated node which features a Haswell processor running at 2.6 GHz with 512 GB of RAM and a 64-bit CentOS Linux 7 operating system. We report average running time in seconds over ten iterations using `ReACt, METAL`, and `ASSET`. In the case of METAL we evaluated the performance of the latest release in GitHub [23]. In each iteration, two or four sets of summary statistics (for 100,000 SNPs) were meta-analyzed. Recall that all methods scale as a function of the number of SNPs and is independent of the number of samples, since only summary statistics are used.

|  | ReACT | METAL | ASSET |
| --- | --- | --- | --- |
| 2 input studies | 2.2s | 1.8s | 696s |
| 4 input studies | 3.1s | 3.3s | 3715s |

**Table S2:**
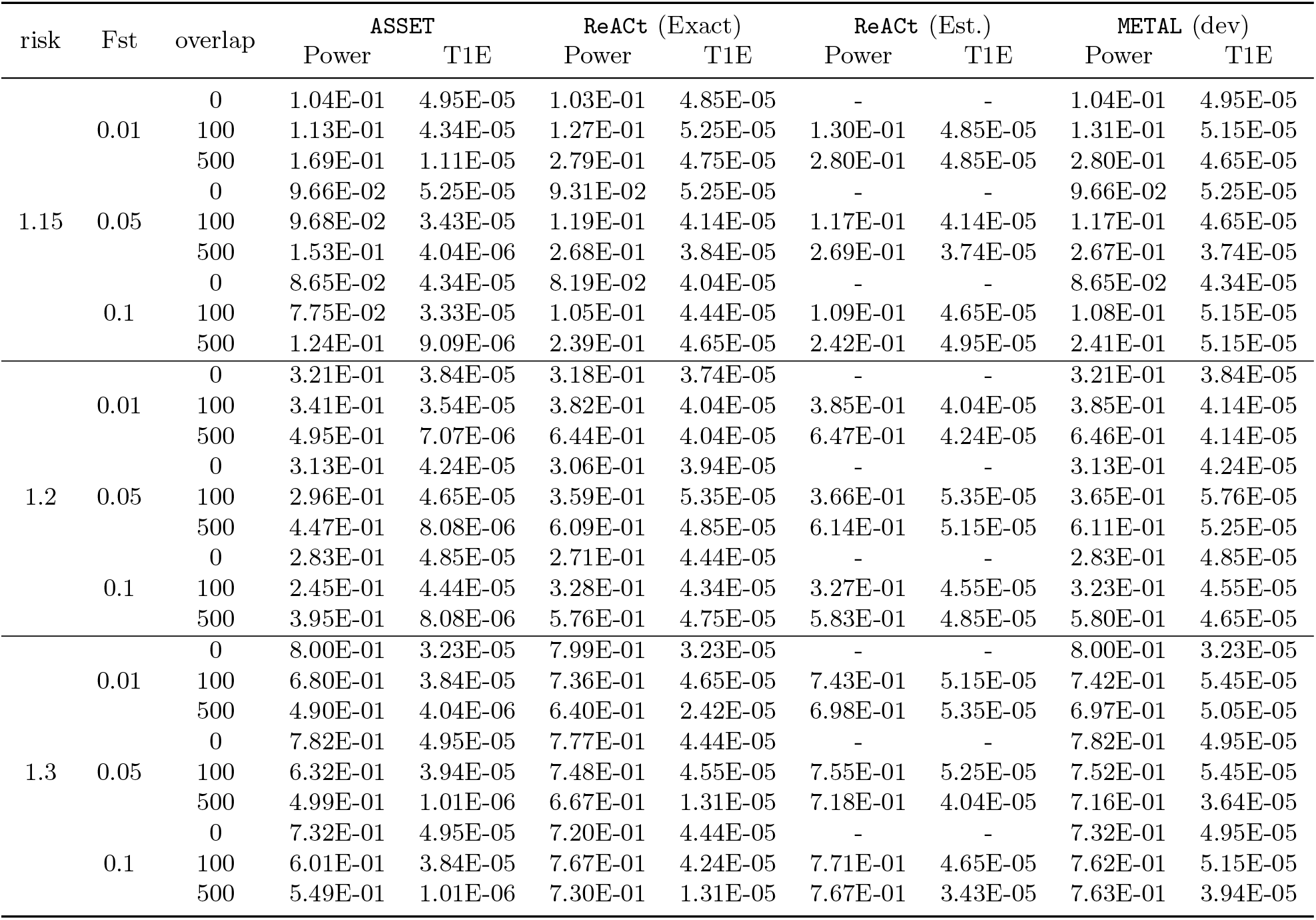
Performance of fixed-effect meta-analysis with two input studies under different conditions. We compare power and type I error rate (T1E) of our method meta-analyzing two studies vs. ASSET/METAL for a significance threshold *p* < 5·10^−5^. METAL dev refers to the latest release in GitHub [23]. Two variants of ReACt are tested: Exact and Est, indicating whether the sample overlap was *exactly* known as part of the input or whether it was *estimated*, respectively. Sample overlap indicates the number of cases and controls that were shared between two input studies. I.e. a sample overlap equal to 100 means that there are 100 cases **and** 100 controls shared between two input studies. Total sample sizes for each input study, including the shared samples, are equal to 2000 when the sample overlap is equal to zero; 2400 when the sample overlap is equal to 100; and 4000 when the sample overlap is equal to 500. In each case, the sample is equally split to cases and controls. Also see figure 1 and 2.

**Table S3:** Performance of fixed-effect meta-analysis with four input studies under different conditions. We compare power and type I error rate (T1E) of our method meta-analyzing four studies vs. ASSET/METAL for a significance threshold *p* < 5·10^−5^. METAL dev refers to the latest release in GitHub [23]. Two variants of ReACt are tested: Exact and Est, indicating whether the sample overlap was *exactly* known as part of the input or whether it was *estimated*, respectively. Sample overlap indicates the number of cases and controls that were shared between two input studies. I.e. a sample overlap equal to 100 means that there are 100 cases **and** 100 controls shared between two input studies. Total sample sizes for each input study, including the shared samples, are equal to 2000 when the sample overlap is equal to zero; 2400 when the sample overlap is equal to 100; and 4000 when the sample overlap is equal to 500. In each case, the sample is equally split to cases and controls.

| risk | Fst | overlap | ASSET |  | ReACt (Exact) |  | ReACt (Est.) |  | METAL (dev) |  |
| --- | --- | --- | --- | --- | --- | --- | --- | --- | --- | --- |
|  |  |  | Power | T1E | Power | T1E | Power | T1E | Power | T1E |
| 1.15 | 0.01 | 0 | 4.31E-01 | 4.75E-05 | 4.31E-01 | 4.75E-05 | - | - | 4.31E-01 | 4.75E-05 |
|  |  | 100 | 3.19E-01 | 2.93E-05 | 4.00E-01 | 5.15E-05 | 4.03E-01 | 5.45E-05 | 4.03E-01 | 4.85E-05 |
|  |  | 500 | 2.36E-01 | 1.01E-06 | 5.20E-01 | 4.85E-05 | 5.27E-01 | 5.25E-05 | 5.23E-01 | 4.85E-05 |
|  | 0.05 | 0 | 4.13E-01 | 4.34E-05 | 4.08E-01 | 4.24E-05 | - | - | 4.13E-01 | 4.34E-05 |
|  |  | 100 | 2.49E-01 | 3.33E-05 | 3.83E-01 | 5.25E-05 | 3.85E-01 | 5.66E-05 | 3.78E-01 | 5.56E-05 |
|  |  | 500 | 2.06E-01 | 2.02E-06 | 5.03E-01 | 5.56E-05 | 5.14E-01 | 6.46E-05 | 5.04E-01 | 5.25E-05 |
|  | 0.1 | 0 | 3.72E-01 | 5.35E-05 | 3.64E-01 | 4.85E-05 | - | - | 3.72E-01 | 5.35E-05 |
|  |  | 100 | 1.90E-01 | 2.42E-05 | 3.46E-01 | 4.55E-05 | 3.53E-01 | 5.66E-05 | 3.41E-01 | 5.45E-05 |
|  |  | 500 | 1.60E-01 | 2.02E-06 | 4.56E-01 | 5.15E-05 | 4.66E-01 | 5.45E-05 | 4.61E-01 | 5.35E-05 |
| 1.2 | 0.01 | 0 | 7.87E-01 | 5.15E-05 | 7.85E-01 | 5.15E-05 | - | - | 7.87E-01 | 5.15E-05 |
|  |  | 100 | 6.48E-01 | 4.14E-05 | 7.59E-01 | 4.85E-05 | 7.64E-01 | 5.45E-05 | 7.59E-01 | 4.95E-05 |
|  |  | 500 | 6.14E-01 | 0.00E+00 | 8.43E-01 | 5.05E-05 | 8.49E-01 | 5.96E-05 | 8.48E-01 | 5.25E-05 |
|  | 0.05 | 0 | 7.61E-01 | 3.43E-05 | 7.57E-01 | 3.23E-05 | - | - | 7.61E-01 | 3.43E-05 |
|  |  | 100 | 5.26E-01 | 1.82E-05 | 7.32E-01 | 3.54E-05 | 7.41E-01 | 4.85E-05 | 7.33E-01 | 4.65E-05 |
|  |  | 500 | 5.36E-01 | 1.01E-06 | 8.19E-01 | 2.93E-05 | 8.28E-01 | 3.54E-05 | 8.23E-01 | 3.23E-05 |
|  | 0.1 | 0 | 7.21E-01 | 5.15E-05 | 7.11E-01 | 5.15E-05 | - | - | 7.21E-01 | 5.15E-05 |
|  |  | 100 | 4.22E-01 | 3.43E-05 | 6.88E-01 | 5.35E-05 | 6.86E-01 | 5.15E-05 | 6.76E-01 | 6.16E-05 |
|  |  | 500 | 4.65E-01 | 1.01E-06 | 7.86E-01 | 4.65E-05 | 7.91E-01 | 5.25E-05 | 7.88E-01 | 5.15E-05 |
| 1.3 | 0.01 | 0 | 9.83E-01 | 5.45E-05 | 9.83E-01 | 5.45E-05 | - | - | 9.83E-01 | 5.45E-05 |
|  |  | 100 | 8.59E-01 | 2.02E-05 | 9.45E-01 | 3.23E-05 | 9.54E-01 | 4.95E-05 | 9.50E-01 | 4.85E-05 |
|  |  | 500 | 6.30E-01 | 0.00E+00 | 8.53E-01 | 5.05E-06 | 9.12E-01 | 6.46E-05 | 9.10E-01 | 6.87E-05 |
|  | 0.05 | 0 | 9.71E-01 | 4.65E-05 | 9.70E-01 | 4.44E-05 | - | - | 9.71E-01 | 4.65E-05 |
|  |  | 100 | 7.68E-01 | 2.22E-05 | 9.49E-01 | 3.23E-05 | 9.55E-01 | 5.15E-05 | 9.50E-01 | 4.85E-05 |
|  |  | 500 | 6.10E-01 | 0.00E+00 | 8.73E-01 | 1.01E-05 | 9.23E-01 | 7.07E-05 | 9.21E-01 | 6.67E-05 |
|  | 0.1 | 0 | 9.54E-01 | 5.66E-05 | 9.52E-01 | 4.65E-05 | - | - | 9.54E-01 | 5.66E-05 |
|  |  | 100 | 6.91E-01 | 2.32E-05 | 9.45E-01 | 4.04E-05 | 9.47E-01 | 4.65E-05 | 9.40E-01 | 5.15E-05 |
|  |  | 500 | 6.21E-01 | 0.00E+00 | 8.93E-01 | 1.01E-05 | 9.27E-01 | 4.04E-05 | 9.24E-01 | 4.55E-05 |

**Table S4:** Performance of sample overlap correction for estimating PRS using our method. Assuming 100 cases and 100 controls shared between base and target studies, we compared the corrected PRS statistics estimated using our method with the real statistics of individual level PRS obtained using PRSice2. Comparison was carried out under various levels of stratification between base and target population (*F*_*st*_ = 0, 0.05, and 0.1) and *p*-value thresholds (denoted by *P*-thres in the table) for SNP selection. For both methods, mean PRS represents the estimated group mean PRS for cases and controls; and *p*-val are the *t*-test *p*-values comparing the resulting PRS distribution in cases and controls. For PRSice2, we computed these statistics for all the samples in the target population, including the samples shared with the base population (denoted by All samples), as well as only for samples that are present exclusively in the target population (denoted by Non-overlapping Samples).

| Fst | $P$-thres | trait | Our method (ReACT) | | PRSice2 | | | |
| --- | --- | --- | --- | --- | --- | --- | --- | --- |
|  |  |  | Corrected statistics |  | All samples |  | Non-overlapping Samples |  |
| | | | mean PRS | $p$-val | mean PRS | $p$-val | mean PRS | $p$-val |
| 0<sup>a</sup> | 0.05 | cases | 0.0003 | 4.07E-05 | 0.0012 | 1.09E-54 | 0.0003 | 3.59E-07 |
|  |  | controls | 0.0000 |  | -0.0009 |  | 0.0000 |  |
|  | 0.005 | cases | 0.0034 | 1.28E-04 | 0.0050 | 6.02E-39 | 0.0034 | 1.20E-04 |
|  |  | controls | 0.0024 |  | 0.0008 |  | 0.0025 |  |
| | $5 \cdot 10^{-4}$ | cases | -0.0030 | 2.44E-01 | -0.0008 | 8.96E-12 | -0.0028 | 1.47E-01 |
|  |  | controls | -0.0041 |  | -0.0063 |  | -0.0040 |  |
| | $5 \cdot 10^{-5}$ | cases | 0.0441 | 7.52E-01 | 0.0471 | 2.31E-02 | 0.0449 | 5.46E-01 |
|  |  | controls | 0.0450 |  | 0.0419 |  | 0.0464 |  |
| 0.05<sup>b</sup> | 0.05 | cases | 0.0000 | 5.57E-54 | 0.0002 | 3.55E-111 | 0.0001 | 8.64E-88 |
|  |  | controls | -0.0005 |  | -0.0007 |  | -0.0006 |  |
|  | 0.005 | cases | 0.0001 | 4.21E-62 | 0.0001 | 5.56E-110 | 0.0000 | 3.30E-91 |
|  |  | controls | -0.0019 |  | -0.0025 |  | -0.0024 |  |
| | $5 \cdot 10^{-4}$ | cases | -0.0063 | 1.51E-50 | -0.0067 | 1.72E-77 | -0.0069 | 3.61E-70 |
|  |  | controls | -0.0112 |  | -0.0124 |  | -0.0124 |  |
| | $5 \cdot 10^{-5}$ | cases | -0.0234 | 4.88E-21 | -0.0229 | 3.21E-32 | -0.0232 | 3.04E-29 |
|  |  | controls | -0.0298 |  | -0.0304 |  | -0.0305 |  |
| 0.1<sup>c</sup> | 0.05 | cases | 0.0001 | 7.32E-35 | 0.0004 | 8.05E-90 | 0.0004 | 7.52E-68 |
|  |  | controls | -0.0003 |  | -0.0004 |  | -0.0003 |  |
|  | 0.005 | cases | 0.0004 | 2.14E-52 | 0.0007 | 8.82E-98 | 0.0006 | 3.03E-79 |
|  |  | controls | -0.0014 |  | -0.0017 |  | -0.0015 |  |
| | $5 \cdot 10^{-4}$ | cases | -0.0048 | 3.74E-41 | -0.0048 | 6.51E-60 | -0.0047 | 1.32E-52 |
|  |  | controls | -0.0091 |  | -0.0100 |  | -0.0096 |  |
| | $5 \cdot 10^{-5}$ | cases | 0.0109 | 6.04E-15 | 0.0087 | 7.62E-22 | 0.0088 | 2.47E-19 |
|  |  | controls | 0.0054 |  | 0.0021 |  | 0.0025 |  |

<sup>a</sup> tested with 550 cases and 550 controls from base and target studies respectively
<sup>b</sup> tested with 1,200 cases and 1,200 controls from base and target studies respectively
<sup>c</sup> tested with 1,200 cases and 1,200 controls from base and target studies respectively

**Table S5. Using ReACt to run cc-GWAS cross eight neuropsychiatric disorders.** We applied our method for cc-GWAS to the summary statistics of eight neuropsychiatric disorders from PGC. Each spreadsheet reports the genomewide significant trait differential regions for a pair of disorders analyzed. For each genomic region, statistics and annotation for the leading SNP are reported.

\***Excel table**.

### 6.2 Solving the non-linear system of equations of Section 2.1

For notational simplicity, let 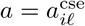, 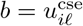, 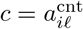 and 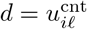. We rewrite eqns. (1)-(4) as

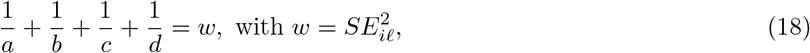

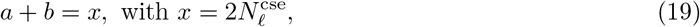

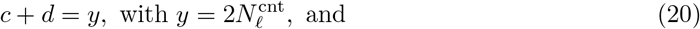

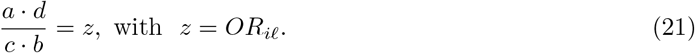

Our goal is compute values for the four unknowns *a*, *b*, *c*, and *d*. Combining eqns. (19) and (20), we get

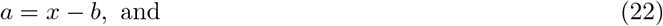

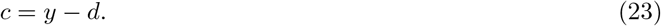

Substituting eqn. (22) and eqn. (23) into eqn. (21), we get (*x − b*)*d* = *zb*(*y − d*), which can be rewritten as

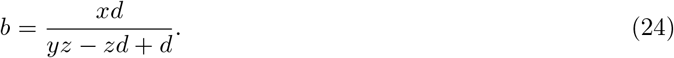

Substituting eqn. (24) into eqn. (22), we get

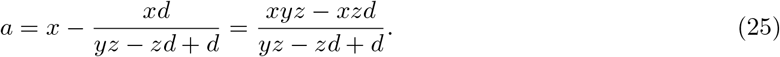

We now note that all four unknowns can be written as functions of *d* and other known quantities. Substituting eqn. (23), eqn. (24), and eqn. (25) into eqn. (18), we get

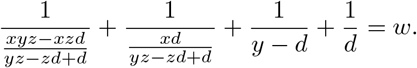

Simplifying the above equation, we get

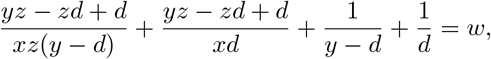

which can be further simplified to

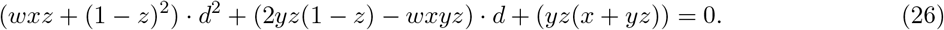

Eqn. (26) is a quadratic equation on *d*; its real roots (if they exist) are

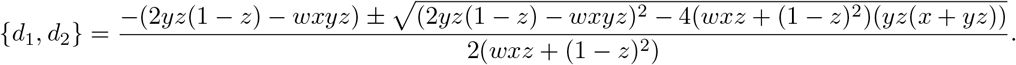

Given *d*, we can immediately compute *a*, *b*, and *c* using eqns. (23), (24), and (25). In order to determine whether *d* is equal to *d*_1_ or *d*_2_, we first check whether *d*_1_ or *d*_2_ guarantee that *a*, *b*, *c*, and *d* are all positive numbers. If both *d*_1_ and *d*_2_ satisfy this constraint, then we choose the *largest* of the two roots, as it solves the following trivial minimization problem:

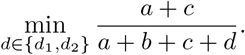

The above choice is based on the assumption that in summary statistics *A*_1_ (whose frequency is equal to the above fraction) typically denotes the affected (minor) allele. Additionally, our code performs a sanity check for allele alignment across studies given the solution *d*_1_ or *d*_2_.

For the sake of completeness, we also prove that it is not possible for both *d*_1_ and *d*_2_ to be negative. First, note that

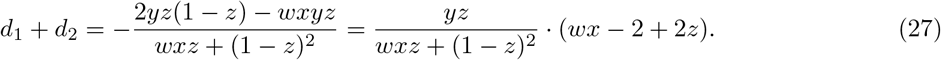

Using *x* = *a* + *b* > 0 and 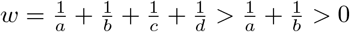, we get

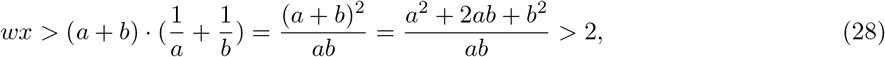

which implies that *wx* − 2 + 2*z* > 0. Combining with eqn. (27), we conclude that *d*_1_ + *d* + 2 is non-negative; recall that *w*, *x*, *y*, and *z* are all non-negative. Additionally,

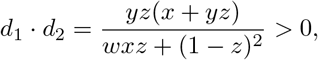

which implies that *d*_1_ and *d*_2_ must have the same sign, and since their sum is non-negative, they must both be positive. It is a simple exercise to prove that as long as root(s) exist, at least one of them will guarantee that all values for *a*, *b* and *c* will be positive.

One important exception arises when the discriminant in eqn. (26) is negative. In that case, no real roots exist for the quadratic equation. We do note that, theoretically, this should never happen, since the underlying unknown quantities are positive real numbers. However, stratification correction and genotype missingness could force the discriminant to fall below zero. To address this issue, we inflate *w* (i.e., the square of the standard error for the respective SNP) and recompute the discriminant. More specifically, we iteratively multiply *w* by 1.001 (a 0.1% inflation) until a non-negative discriminant is obtained or until 50 iterations are reached. The maximum inflation we allow (after the full 50 iterations) is 1.001^50^ − 1 ≈ 5%. If after 50 iterations we have failed to find a non-negative discriminant we omit this particular SNP from further analyses. Empirically, for most input SNPs, a real root can be found after at most ten iterations.

### 6.3 Correction for sample overlap between the base/target studies for group PRS

The existence of shared samples in base (discovery) and target populations can lead to inflation in association between PRS and the trait of interest in the target population [46, 19]. In our case, such overlap will cause higher levels of significance in the *t*-test comparing the case and control PRS distribution. So far, for conventional PRS, the most widely accepted approach to address this problem is simply to identify the overlapping individuals and remove them from the target population. However, in practice, this is not always possible since it usually requires additional access to the individual level data of the base population. In this section, we introduce a correction for sample overlap between the base and target populations implemented in ReACt that could alleviate such issues.

In the following, we will use the case group as an example. Assume that the sample size for cases of the target population is 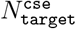, out of which 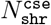 are also cases in the base population (overlap). If the probability of each sample being shared between the base and target studies is uniformly distributed in both base and target studies, we would expect the observed mean PRS in target cases 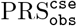 to be a weighted sum of the mean PRS in base cases 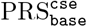 and the mean PRS of cases that only exist in the target population 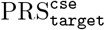 as follows:

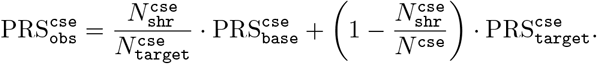

Therefore, the mean PRS for cases only in the target population is:

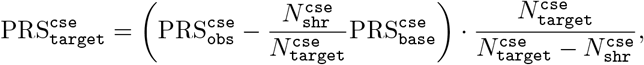

where 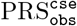 is the uncorrected group mean computed as described in Section 4.3.1. 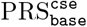 can be obtained by simply setting the target population to be the same as the base population, using base summary
statistics to compute group PRS for the target population. Similarly, we can adjust the variance computation as follows:

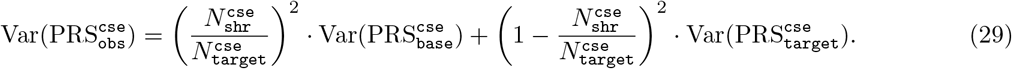

Therefore, the corrected variance will be

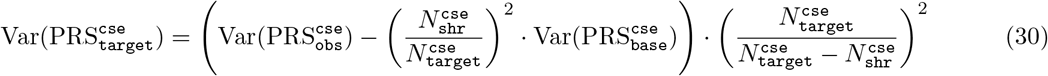

Similarly,

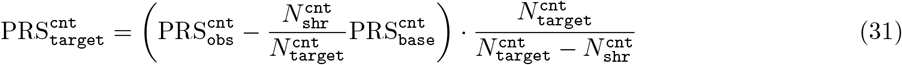

and

s
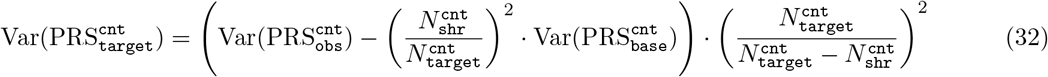

for controls. Then, the corrected *p*-value will be based on a *t*-test using the corrected mean and variance and an adjusted degree of freedom:

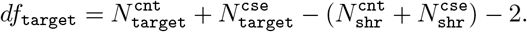

This is a straightforward correction on the target PRS using the scores of the base population that one would use if there were no stratification between the base and target populations. In practice, this idealized scenario does not hold. In order to deal with the stratification between the base and target populations, prior to any correction, we shift the scores for base cases and controls by aligning the base population PRS means to the target population as follows:

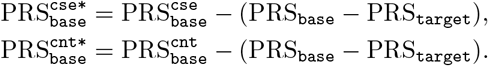

In the above, PRS_base_ and PRS_target_ are mean PRS for the base and target populations respectively:

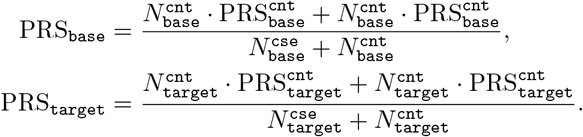

In practice, we use 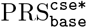 and 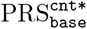 instead of 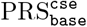 and 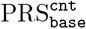 in equations (29)-(32) for correction. We evaluated the performance of this correction scheme by introducing sample overlaps between the base and target populations using the same simulation model as the one we used to evaluate the performance of our group PRS approach. We computed the real individual level PRS using PRSice2, from which we obtained the inflated PRS descriptive statistics (group mean, standard deviation, and *t*-test *p*-value) for all target samples, including the ones that are shared with the base population. We also computed PRS statistics for samples that are present only in the target population as the ground truth. We compared results from our corrected group PRS method to the PRS statistics for the samples that are exclusive to the target population computed using PRSice2. Results on synthetic data demonstrated that our correction can drastically alleviate the inflation in *p*-values that is the result of sample overlap the between base and target populations. See Table S4, which shows representative results from our experimental evaluations. If the number of overlapping samples is unknown to the user, we apply the approach proposed in [23] to get an estimate of the overlapping sample size and we correct the output statistics accordingly. Note that this correction approach is based on the assumption that all samples having an equal probability of being shared between the base and target populations, which might be unrealistic in certain settings.

## Footnotes

1 This is simply the sum of the absolute values of the three entries of the vector *β*^t+1^ − *β*^t^.

## References

[1] David W Craig, Robert M Goor, Zhenyuan Wang, Justin Paschall, Jim Ostell, Michael Feolo, Stephen T Sherry, and Teri A Manolio. Assessing and managing risk when sharing aggregate genetic variant data. Nature Reviews Genetics, 12(10):730–736, 2011.

[2] Bogdan Pasaniuc and Alkes L Price. Dissecting the genetics of complex traits using summary association statistics. Nature Reviews Genetics, 18(2):117, 2017.

[3] Ju-Hyun Park, Sholom Wacholder, Mitchell H Gail, Ulrike Peters, Kevin B Jacobs, Stephen J Chanock, and Nilanjan Chatterjee. Estimation of effect size distribution from genome-wide association studies and implications for future discoveries. Nature genetics, 42(7):570–575, 2010.

[4] Yan Zhang, Guanghao Qi, Ju-Hyun Park, and Nilanjan Chatterjee. Estimation of complex effect-size distributions using summary-level statistics from genome-wide association studies across 32 complex traits. Nature genetics, 50(9):1318–1326, 2018.

[5] Zhiyu Yang, Hanrui Wu, Phil H Lee, Fotis Tsetsos, Lea K Davis, Dongmei Yu, Sang Hong Lee, Søren Dalsgaard, Jan Haavik, Csaba Barta, et al. Investigating shared genetic basis across tourette syndrome and comorbid neurodevelopmental disorders along the impulsivity-compulsivity spectrum. Biological Psychiatry.

[6] Fotis Tsetsos, Shanmukha S Padmanabhuni, John Alexander, Iordanis Karagiannidis, Margaritis Tsifintaris, Apostolia Topaloudi, Dimitrios Mantzaris, Marianthi Georgitsi, Petros Drineas, and Peristera Paschou. Meta-analysis of tourette syndrome and attention deficit hyperactivity disorder provides support for a shared genetic basis. Frontiers in neuroscience, 10:340, 2016.

[7] Phil H Lee, Verneri Anttila, Hyejung Won, Yen-Chen A Feng, Jacob Rosenthal, Zhaozhong Zhu, Elliot M Tucker-Drob, Michel G Nivard, Andrew D Grotzinger, Danielle Posthuma, et al. Genomic relationships, novel loci, and pleiotropic mechanisms across eight psychiatric disorders. Cell, 179(7):1469–1482, 2019.

[8] Daniel J Schaid, Wenan Chen, and Nicholas B Larson. From genome-wide associations to candidate causal variants by statistical fine-mapping. Nature Reviews Genetics, 19(8):491–504, 2018.

[9] Christian Benner, Chris CA Spencer, Aki S Havulinna, Veikko Salomaa, Samuli Ripatti, and Matti Pirinen. Finemap: efficient variable selection using summary data from genome-wide association studies. Bioinformatics, 32(10):1493–1501, 2016.

[10] Bogdan Pasaniuc, Noah Zaitlen, Huwenbo Shi, Gaurav Bhatia, Alexander Gusev, Joseph Pickrell, Joel Hirschhorn, David P Strachan, Nick Patterson, and Alkes L Price. Fast and accurate imputation of summary statistics enhances evidence of functional enrichment. Bioinformatics, 30(20):2906–2914, 2014.

[11] Sina Rüeger, Aaron McDaid, and Zoltán Kutalik. Evaluation and application of summary statistic imputation to discover new height-associated loci. PLoS genetics, 14(5):e1007371, 2018.

[12] Brendan K Bulik-Sullivan, Po-Ru Loh, Hilary K Finucane, Stephan Ripke, Jian Yang, Nick Patterson, Mark J Daly, Alkes L Price, and Benjamin M Neale. Ld score regression distinguishes confounding from polygenicity in genome-wide association studies. Nature genetics, 47(3):291–295, 2015.

[13] Brielin C Brown, Chun Jimmie Ye, Alkes L Price, Noah Zaitlen, Asian Genetic Epidemiology Network Type 2 Diabetes Consortium, et al. Transethnic genetic-correlation estimates from summary statistics. The American Journal of Human Genetics, 99(1):76–88, 2016.

[14] Jie Zheng, A Mesut Erzurumluoglu, Benjamin L Elsworth, John P Kemp, Laurence Howe, Philip C Haycock, Gibran Hemani, Katherine Tansey, Charles Laurin, Beate St Pourcain, et al. Ld hub: a centralized database and web interface to perform ld score regression that maximizes the potential of summary level gwas data for snp heritability and genetic correlation analysis. Bioinformatics, 33(2):272– 279, 2017.

[15] Hilary K Finucane, Brendan Bulik-Sullivan, Alexander Gusev, Gosia Trynka, Yakir Reshef, Po-Ru Loh, Verneri Anttila, Han Xu, Chongzhi Zang, Kyle Farh, et al. Partitioning heritability by functional annotation using genome-wide association summary statistics. Nature genetics, 47(11):1228, 2015.

[16] Wouter J Peyrot and Alkes L Price. Identifying loci with different allele frequencies among cases of eight psychiatric disorders using cc-gwas. Nature Genetics, pages 1–10, 2021.

[17] Robert A Power, Stacy Steinberg, Gyda Bjornsdottir, Cornelius A Rietveld, Abdel Abdellaoui, Michel M Nivard, Magnus Johannesson, Tessel E Galesloot, Jouke J Hottenga, Gonneke Willemsen, et al. Polygenic risk scores for schizophrenia and bipolar disorder predict creativity. Nature neuroscience, 18(7):953–955, 2015.

[18] Ali Torkamani, Nathan E Wineinger, and Eric J Topol. The personal and clinical utility of polygenic risk scores. Nature Reviews Genetics, 19(9):581–590, 2018.

[19] Shing Wan Choi, Timothy Shin-Heng Mak, and Paul F O’Reilly. Tutorial: a guide to performing polygenic risk score analyses. Nature Protocols, 15(9):2759–2772, 2020.

[20] Andrew D Grotzinger, Mijke Rhemtulla, Ronald de Vlaming, Stuart J Ritchie, Travis T Mallard, W David Hill, Hill F Ip, Riccardo E Marioni, Andrew M McIntosh, Ian J Deary, et al. Genomic structural equation modelling provides insights into the multivariate genetic architecture of complex traits. Nature human behaviour, 3(5):513–525, 2019.

[21] Cristen J Willer, Yun Li, and Gonçalo R Abecasis. Metal: fast and efficient meta-analysis of genomewide association scans. Bioinformatics, 26(17):2190–2191, 2010.

[22] Samsiddhi Bhattacharjee, Preetha Rajaraman, Kevin B Jacobs, William A Wheeler, Beatrice S Melin, Patricia Hartge, Meredith Yeager, Charles C Chung, Stephen J Chanock, Nilanjan Chatterjee, et al. A subset-based approach improves power and interpretation for the combined analysis of genetic as-sociation studies of heterogeneous traits. The American Journal of Human Genetics, 90(5):821–835, 2012.

[23] Sebanti Sengupta. Metal, unpublished material and methods. https://genome.sph.umich.edu/w/images/7/7b/METAL_sample_overlap_method_2017-11-15.pdf.

[24] Shing Wan Choi and Paul F O’Reilly. Prsice-2: Polygenic risk score software for biobank-scale data. Gigascience, 8(7):giz082, 2019.

[25] Apostolia Topaloudi, Zoi Zagoriti, Alyssa C Flint, Melanie B Martinez, Zhiyu Yang, Fotis Tsetsos, Yiolanda-Panayiota Christou, George Lagoumintzis, Evangelia Yannaki, Eleni Papanicolaou-Zamba, et al. A myasthenia gravis genomewide association study of three cohorts identifies agrin as a novel risk locus. medRxiv, 2020.

[26] Eli A Stahl, Gerome Breen, Andreas J Forstner, Andrew McQuillin, Stephan Ripke, Vassily Trubetskoy, Manuel Mattheisen, Yunpeng Wang, Jonathan RI Coleman, Héléna A Gaspar, et al. Genome-wide association study identifies 30 loci associated with bipolar disorder. Nature genetics, 51(5):793–803, 2019.

[27] Schizophrenia Working Group of the Psychiatric Genomics Consortium et al. Biological insights from 108 schizophrenia-associated genetic loci. Nature, 511(7510):421–427, 2014.

[28] Douglas M Ruderfer, Stephan Ripke, Andrew McQuillin, James Boocock, Eli A Stahl, Jennifer M Whitehead Pavlides, Niamh Mullins, Alexander W Charney, Anil PS Ori, Loes M Olde Loohuis, et al. Genomic dissection of bipolar disorder and schizophrenia, including 28 subphenotypes. Cell, 173(7):1705–1715, 2018.

[29] W Scott Watkins, Alan R Rogers, Christopher T Ostler, Steve Wooding, Michael J Bamshad, Anna-Marie E Brassington, Marion L Carroll, Son V Nguyen, Jerilyn A Walker, BV Ravi Prasad, et al. Genetic variation among world populations: inferences from 100 alu insertion polymorphisms. Genome research, 13(7):1607–1618, 2003.

[30] L Duncan, H Shen, B Gelaye, J Meijsen, K Ressler, M Feldman, R Peterson, and B Domingue. Analysis of polygenic risk score usage and performance in diverse human populations. Nature communications, 10(1):1–9, 2019.

[31] Frank Dudbridge. Power and predictive accuracy of polygenic risk scores. PLoS Genet, 9(3):e1003348, 2013.

[32] Logistic regression. http://nlp.chonbuk.ac.kr/BML/slides_freda/lec7.pdf. Accessed: 2020-04-13.

[33] Christopher C Chang, Carson C Chow, Laurent CAM Tellier, Shashaank Vattikuti, Shaun M Pur-cell, and James J Lee. Second-generation plink: rising to the challenge of larger and richer datasets. Gigascience, 4(1):s13742–015, 2015.

[34] David Firth. Bias reduction of maximum likelihood estimates. Biometrika, pages 27–38, 1993.

[35] Georg Heinze and Michael Schemper. A solution to the problem of separation in logistic regression. Statistics in medicine, 21(16):2409–2419, 2002.

[36] Clement Ma, Tom Blackwell, Michael Boehnke, Laura J Scott, and GoT2D Investigators. Recommended joint and meta-analysis strategies for case-control association testing of single low-count variants. Ge-netic epidemiology, 37(6):539–550, 2013.

[37] Pedro RD Bom and Heiko Rachinger. A generalized-weights solution to sample overlap in meta-analysis. Research Synthesis Methods, 2020.

[38] Paul D Arnold, Kathleen D Askland, Cristina Barlassina, Laura Bellodi, OJ Bienvenu, Donald Black, Michael Bloch, Helena Brentani, Christie L Burton, Beatriz Camarena, et al. Revealing the com-plex genetic architecture of obsessive-compulsive disorder using meta-analysis. Molecular psychiatry, 23(5):1181–1181, 2018.

[39] Dongmei Yu, Jae Hoon Sul, Fotis Tsetsos, Muhammad S Nawaz, Alden Y Huang, Ivette Zelaya, Cornelia Illmann, Lisa Osiecki, Sabrina M Darrow, Matthew E Hirschtritt, et al. Interrogating the genetic determinants of tourette’s syndrome and other tic disorders through genome-wide association studies. American Journal of Psychiatry, 176(3):217–227, 2019.

[40] Laramie Duncan, Zeynep Yilmaz, Helena Gaspar, Raymond Walters, Jackie Goldstein, Verneri Anttila, Brendan Bulik-Sullivan, Stephan Ripke, Eating Disorders Working Group of the Psychiatric Genomics Consortium, Laura Thornton, et al. Significant locus and metabolic genetic correlations revealed in genome-wide association study of anorexia nervosa. American journal of psychiatry, 174(9):850–858, 2017.

[41] Jakob Grove, Stephan Ripke, Thomas D Als, Manuel Mattheisen, Raymond K Walters, Hyejung Won, Jonatan Pallesen, Esben Agerbo, Ole A Andreassen, Richard Anney, et al. Identification of common genetic risk variants for autism spectrum disorder. Nature genetics, 51(3):431–444, 2019.

[42] Ditte Demontis, Raymond K Walters, Joanna Martin, Manuel Mattheisen, Thomas D Als, Esben Agerbo, Gísli Baldursson, Rich Belliveau, Jonas Bybjerg-Grauholm, Marie Bækvad-Hansen, et al. Dis-covery of the first genome-wide significant risk loci for attention deficit/hyperactivity disorder. Nature genetics, 51(1):63–75, 2019.

[43] Naomi R Wray, Stephan Ripke, Manuel Mattheisen, Maciej Trzaskowski, Enda M Byrne, Abdel Abdel-laoui, Mark J Adams, Esben Agerbo, Tracy M Air, Till MF Andlauer, et al. Genome-wide association analyses identify 44 risk variants and refine the genetic architecture of major depression. Nature genetics, 50(5):668–681, 2018.

[44] Schizophrenia Working Group of the Psychiatric Genomics Consortium et al. Biological insights from 108 schizophrenia-associated genetic loci. Nature, 511(7510):421–427, 2014.

[45] Eli A Stahl, Gerome Breen, Andreas J Forstner, Andrew McQuillin, Stephan Ripke, Vassily Trubetskoy, Manuel Mattheisen, Yunpeng Wang, Jonathan RI Coleman, Héléna A Gaspar, et al. Genome-wide association study identifies 30 loci associated with bipolar disorder. Nature genetics, 51(5):793–803, 2019.

[46] Naomi R Wray, Jian Yang, Ben J Hayes, Alkes L Price Michael E Goddard, and Peter M Visscher. Pitfalls of predicting complex traits from snps. Nature Reviews Genetics, 14(7):507–515, 2013.

